# FUNDAMENTAL LIMITATIONS OF KILOHERTZ-FREQUENCY CARRIERS IN AFFERENT FIBER RECRUITMENT WITH TRANSCUTANEOUS SPINAL CORD STIMULATION

**DOI:** 10.1101/2024.07.26.603982

**Authors:** Rodolfo Keesey, Ursula Hofstoetter, Zhaoshun Hu, Lorenzo Lombardi, Rachel Hawthorn, Noah Bryson, Andreas Rowald, Karen Minassian, Ismael Seáñez

## Abstract

The use of kilohertz-frequency (KHF) waveforms has rapidly gained momentum in transcutaneous spinal cord stimulation (tSCS) to restore motor function after paralysis. However, the mechanisms by which these fast-alternating currents depolarize efferent and afferent fibers remain unknown. Our study fills this research gap by providing a hypothesis-and evidence-based investigation using peripheral nerve stimulation, lumbar tSCS, and cervical tSCS in 25 unimpaired participants together with computational modeling. Peripheral nerve stimulation experiments and computational modeling showed that KHF waveforms negatively impact the processes required to elicit action potentials, thereby increasing response thresholds and biasing the recruitment towards efferent fibers. While these results translate to tSCS, we also demonstrate that lumbar tSCS results in the preferential recruitment of afferent fibers, while cervical tSCS favors recruitment of efferent fibers. Given the assumed importance of proprioceptive afferents in motor recovery, our work suggests that the use of KHF waveforms should be reconsidered to maximize neurorehabilitation outcomes, particularly for cervical tSCS. We posit that careful analysis of the mechanisms that mediate responses elicited by novel approaches in tSCS is crucial to understanding their potential to restore motor function after paralysis.

## Main

Spinal cord injury (SCI) is a life-altering event that leads to long-lasting motor impairment, and currently, there is no cure for paralysis^1^. Recent work combining invasive or non-invasive spinal cord stimulation with activity-based training has shown unprecedented improvements in motor function in the chronic stage of paralysis (>1-year post-injury)^2–7^. Such findings have rapidly accelerated the clinical, research, and commercial interest in spinal cord stimulation, promoting the exploration of novel approaches to maximize recovery outcomes. New research directions, including innovative electrode configurations and stimulation patterns, reflect a broader initiative to enhance and optimize recovery in individuals with SCI ^8–10^. However, the specific neural substrates targeted by these novel techniques and the mechanisms by which they are activated remain largely understudied. In the present study, we use human neurophysiology and computational modeling to explore these fundamental questions on two common stimulation waveforms currently employed in transcutaneous spinal cord stimulation (tSCS).

Decades of research in computational modeling ^6,11,12^ and neurophysiological studies in animals^13,14^ and humans^15,16^ support the notion that the motor effects evoked by epidural spinal cord stimulation (eSCS) are elicited by artificially induced action potentials in medium to large-diameter somatosensory fibers within the posterior roots^17,18^. Therefore, the activity of specific neural networks within the spinal cord can be effectively modulated by electrical stimulation of their afferents—a fundamental aspect of neuromodulation^19^. Moreover, activation provided via afferent fibers enables interaction with supraspinal and spinal inputs at the spinal circuit level^14,20,21^, which is thought to be necessary to promote neuroplasticity and functional recovery^22–25^.

In conventional applications of tSCS, biphasic pulses with 0.5-2 ms widths are delivered at 30-50 Hz to facilitate motor function^26–30^. However, the intense cutaneous and neuromuscular co-activation beneath the stimulating surface electrodes has led to the adoption of kilohertz-frequency (KHF) waveforms, typically modulating 0.5-1 ms pulses with a 5-10 kHz carrier^31–33^. These KHF waveforms were originally used in neuromuscular stimulation and were claimed to maximize generated limb torques at tolerable stimulation intensities^34,35^. However, while conventional waveforms in lumbar tSCS have been shown to activate the posterior roots via similar mechanisms as eSCS ^16,36–39^, neither the neural substrates targeted by KHF waveforms nor the mechanisms by which they are depolarized have been previously described. Additionally, KHF waveforms have been recently reported to require considerably higher stimulation intensities to elicit muscle responses than those of conventional waveforms^40,41^. Nevertheless, recent clinical studies to improve motor function via tSCS-assisted rehabilitation in people with SCI have already started to adopt KHF waveforms^7,32,42^.

For almost 90 years, research has shown that varying the duration of conventional rectangular pulses differently affects the recruitment thresholds of motor and proprioceptive fibers^43–45^. The strength-duration time constants for proprioceptive fibers have been found to be longer than those for motor fibers, meaning that their membrane potential changes at a slower rate in response to applied electric currents^46,47^. These differences result in longer pulses (∼1 ms) preferentially recruiting proprioceptive fibers and shorter pulses (<200 µs) favoring direct recruitment of motor fibers^43,44,48^. KHF waveforms of 1 ms duration with 10 kHz carriers essentially contain ten, 100 µs, biphasic pulses. Based on the brief, 50 µs depolarization phases, we hypothesized an increased recruitment of motor efferent fibers over proprioceptive afferents by KHF waveforms. We suspected that the alternating positive and negative phases in KHF waveforms would rapidly de-and hyper-polarize the axon membranes. Therefore, for action potentials to be generated by KHF waveforms, the hyperpolarization introduced by the negative phases must be overcome by the aggregation of the depolarization during the positive phases^49^.

To examine these hypotheses, we recorded muscle responses evoked by stimulation of the mixed tibial nerve as a simplified system to understand the recruitment characteristics of non-synaptic direct (motor efferent) and trans-synaptic reflex-mediated (proprioceptive afferent) responses. We analyzed membrane potentials in a computational model recreating this setup with unprecedented anatomical realism. We found that KHF waveforms resulted in the preferential elicitation of motor-mediated responses over reflex-mediated responses due to differences in membrane properties between efferent and afferent fibers. Moreover, we found evidence of summation processes manifesting in increasing response latencies and decreasing threshold stimulation amplitudes for longer KHF waveforms. The computational model revealed that the hyperpolarizing phases of KHF pulses counteract the depolarization of both afferent and efferent fibers, thus interrupting the summation processes that lead to action potentials, which explain differences in threshold amplitudes and latencies between waveforms.

We next sought to understand whether this reduced efficacy of KHF waveforms to recruit afferent fibers translates into the application of lumbar and cervical tSCS. In lumbar tSCS, post-activation depression and longer response latencies compared to peripheral conduction times confirmed activation of proprioceptive afferents within the posterior roots. In contrast, the absence of post-activation depression in cervical tSCS and an equal, or even shorter response latency compared to peripheral conduction time suggests direct activation of the motor efferents. In all metrics, KHF waveforms proved less effective at recruiting the afferent pathways. The increased recruitment of motor fibers by KHF waveforms, bypassing spinal circuits, could be an unintended side effect that undermines the neuromodulatory potential^13,14,19,24^ of tSCS, particularly in the cervical spinal cord. We posit that the adoption of novel tSCS paradigms requires careful analysis of the mechanisms by which they recruit neural structures prior to clinical translation, as these can have important implications in their potential to promote functional recovery.

## Results

To study the recruitment mechanisms of KHF and conventional waveforms, we first conducted stimulation of the mixed tibial nerve (**Fig. 1a** and **Supplementary Fig. 1**). The tibial nerve presents an ideal model for testing the relative efficacy of afferent vs. efferent stimulation, as the fibers elicit distinct EMG potentials based on differences in their conduction times, namely the Hoffman reflex (H-reflex) and the M-wave, respectively^50,51^. In addition, we recreated our experimental setup in-silico using an anatomically accurate volume conductor model of the lower limb incorporating the tibial nerve, in which biophysically distinct afferent and efferent fibers were generated with the same fiber diameter distribution and topology (**Fig. 1b**).

**Figure 1.**
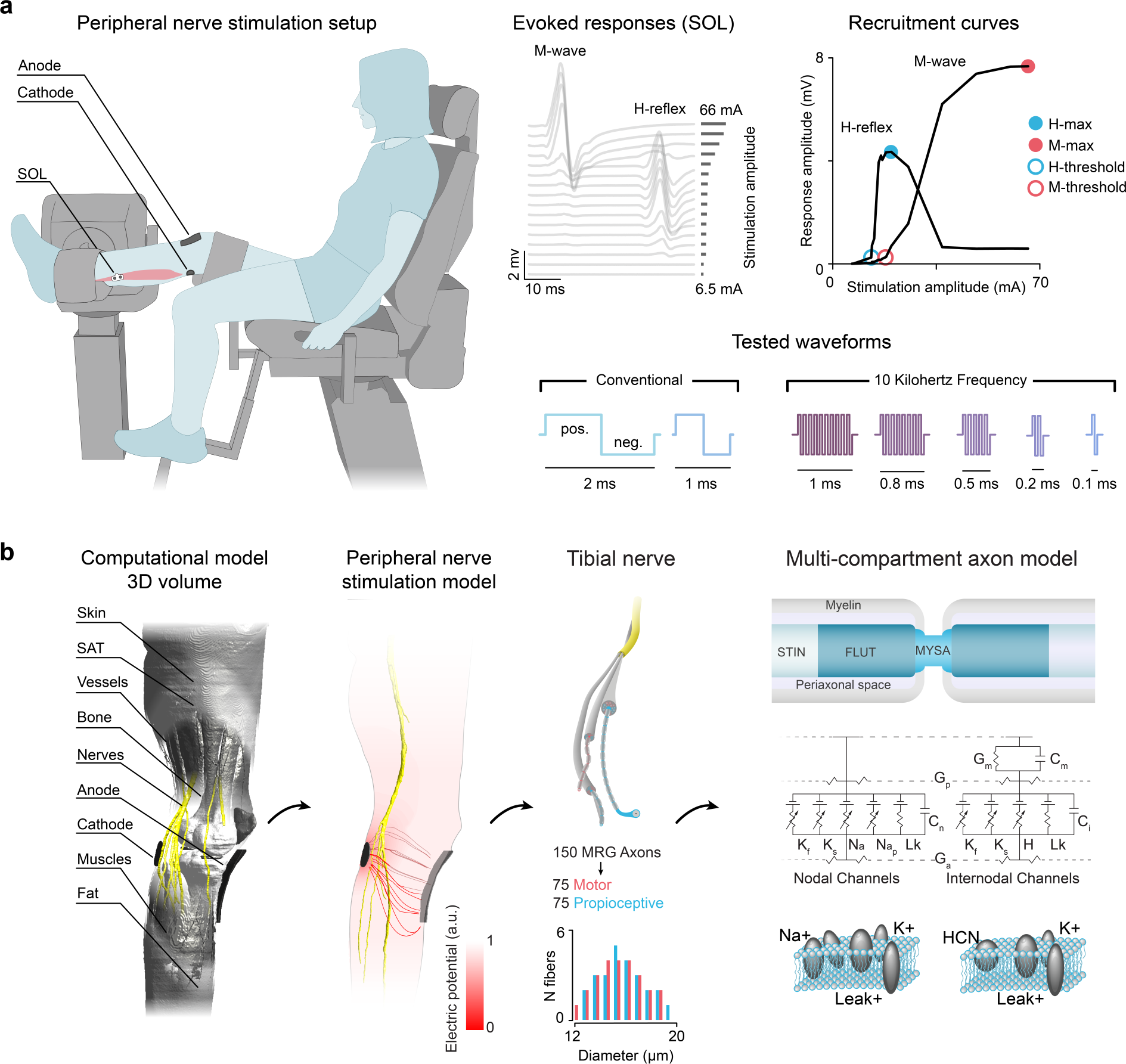
Human neurophysiology experiments and computational modeling in peripheral nerve stimulation to study recruitment mechanisms of conventional and KHF waveforms. **a.** Schematic of the experimental paradigm for peripheral nerve stimulation. We measured EMG activity from the right soleus as participants’ legs were secured in a Biodex System 4 Pro^TM^ isokinetic dynamometer (Biodex Medical Systems, Shirley, NY, USA). We delivered electrical stimulation across the tibial nerve by positioning a cathode at the popliteal fossa and an anode over the lower patella. Threshold stimulation currents and maximum evoked response amplitudes of M-waves and H-reflexes were used to study the recruitment mechanisms of conventional and KHF waveforms in motor and reflex-mediated responses. The positive and negative phases referred to throughout the manuscript are denoted in the conventional waveform. Evoked responses and recruitment curves are shown for participant RS008. **b.** An anatomically accurate 3D volume conductor model of the lower leg was created from the Jeduk model^52–55^ in Sim4Life to replicate the experimental setup from **a.** The electric field distribution was calculated and coupled with neural axon models^56^ of 75 motor efferent and 75 proprioceptive afferent fibers with diameters sampled from the same distribution ranging from 12 to 20 µm^6^. Simulations were performed with the same waveforms that were used in the neurophysiology experiments. Abbreviations: soleus (SOL), positive (pos.), negative (neg.), subcutaneous fat (SAT), stereotyped internodal region (STIN), fluted paranodal main region (FLUT), myelin sheath attachment region (MYSA), hyperpolarization-activated cyclic nucleotide-gated (HCN).

### Kilohertz-frequency waveforms require higher stimulation intensities than conventional waveforms to evoke responses and show evidence of interrupted sub-threshold summation processes

We first investigated the effect of stimulation waveform (conventional biphasic pulses of 1 ms and 2 ms duration as well as KHF bursts at 10 kHz of 0.1 ms, 0.2 ms, 0.5 ms, 0.8 ms and 1.0 ms duration) on the elicitation of H-reflexes and M-waves (**Fig. 2a**). Due to the long-duration depolarization phase (positive phase), we hypothesized that conventional waveforms would require lower stimulation intensities to activate afferent and efferent fibers than any of the KHF waveforms tested. Indeed, statistical analyses demonstrated a significant effect of stimulation waveform on the threshold stimulation currents for eliciting H-reflexes and M-waves (**Supplementary Table 1**). Both the 2 ms and 1 ms conventional waveforms had significantly lower H-reflex and M-wave thresholds than any of the KHF waveforms (**Fig. 2c**). The large discrepancy in thresholds even after accounting for delivered charge (Supplementary Fig. 2, **Supplementary Table 1**) implies different underlying recruitment mechanisms between conventional and KHF waveforms.

**Figure 2.**
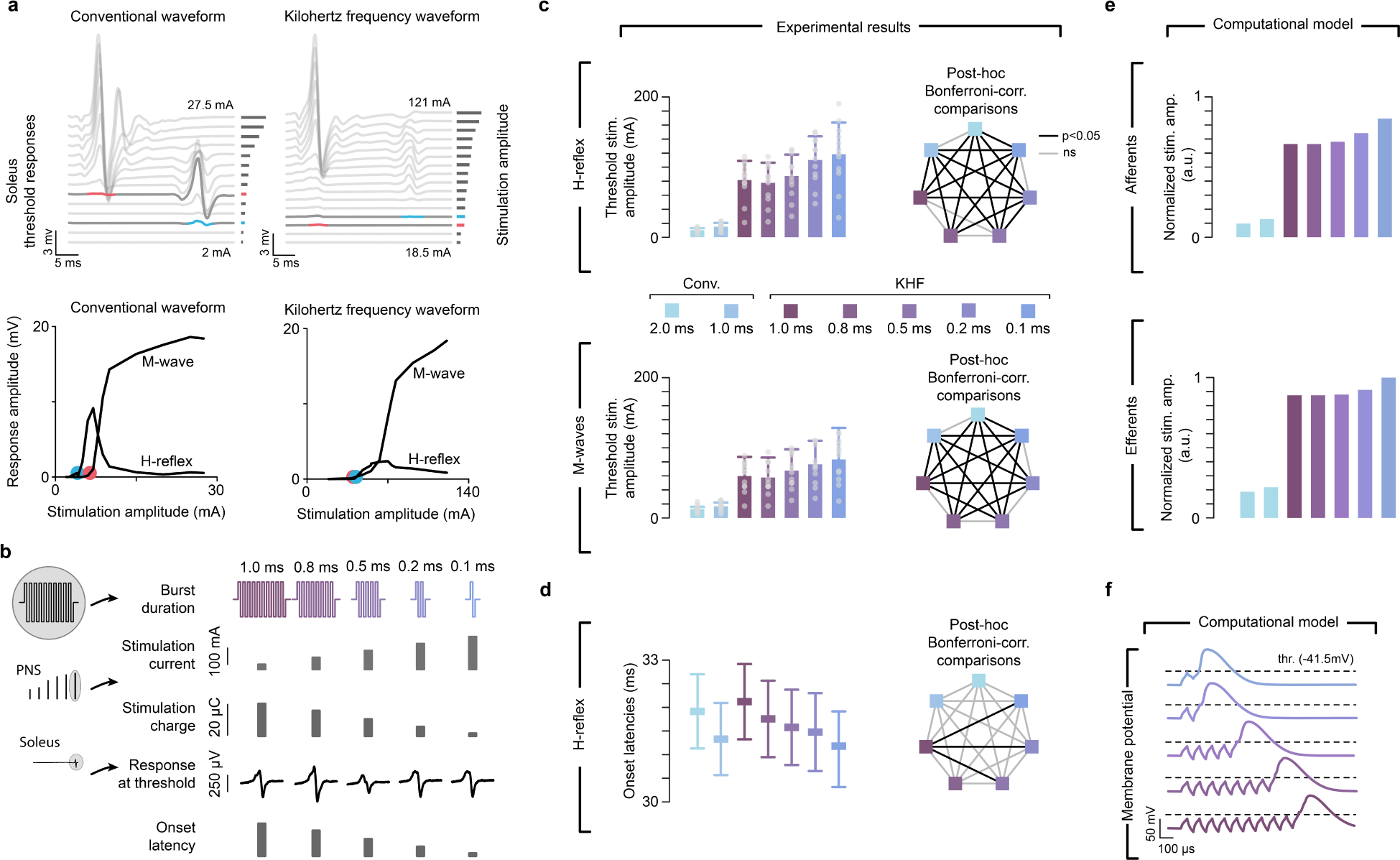
Effect of waveform and duration on threshold stimulation currents and response-latencies. **a.** Soleus responses at increasing stimulation currents with conventional and KHF waveforms in participant RS012. H-reflex and M-wave threshold stimulation currents are defined as the lowest currents eliciting responses with peak-to-peak amplitudes larger than 50 µV. Threshold responses are illustrated in blue for H-reflexes and red for M-waves. The recruitment curves are shown below their respective response traces. The threshold amplitudes are denoted with blue and red circles. **b.** Hypothesis: due to summation processes, increasing burst duration in KHF waveforms decreases the current and increases the charge necessary to evoke responses at threshold as well as increases the H-reflex latencies. Representative stimulation current, charge, and evoked responses are shown for RS008. Onset latencies are illustrations of the expected trend, and not actual data. **c.** Threshold stimulation currents required to evoke responses at threshold decreased with increasing KHF waveform duration; this is true for both the H-reflex and M-wave. **d**. H-reflex response latencies at threshold increased with increasing burst duration. **e**. Threshold stimulation currents for KHF waveforms of different durations. Mirroring our physiological results, our computational model revealed that short KHF waveform durations require higher stimulation amplitudes to recruit both afferent and efferent fibers. **f**. Membrane potentials at threshold amplitudes for KHF waveforms of different burst durations. As the action potential is generated by the last phase of the KHF waveform, response latency decreases with decreasing burst duration (*cf.* **Supplementary Fig. 3**). Bars in **c** represent estimated means while whiskers indicate the standard errors. Central squares in **d** represent the estimated means while whiskers indicate the 95% confidence intervals. The heptagon indicates the results for the post-hoc Bonferroni-corrected comparisons between waveforms; solid black lines indicate statistically significant differences between waveforms (p < 0.05). Abbreviations: kilohertz frequency (KHF), conventional (Conv.), stimulation (stim.), corrected (corr.), amplitude (amp.), arbitrary units (a.u.).

Interestingly, as duration of KHF waveforms increased, the stimulation current required to evoke responses at threshold decreased for both H-reflexes and M-waves (**Fig. 2c**, **Supplementary Table 1**) while required charge increased (**Supplementary Fig. 2**, **Supplementary Table 1).** This finding suggests the presence of summation processes, where the fast alternation in active depolarization and hyperpolarization introduced by the alternating currents in KHF waveforms must be overcome by the aggregation of the depolarization during the positive phases to reach threshold. Therefore, all sub-pulses in the waveform should contribute to evoking threshold responses, increasing the latency of the response (**Fig. 2b**). To test this hypothesis, we compared H-reflex response latencies at 1-1.5 times the threshold. We identified waveform as a significant factor (**Supplementary Table 2**). Specifically, response latencies increased with increasing KHF waveform duration, with a 1.0 ms difference between 1.0 ms and 0.1 ms KHF waveforms (**Fig. 2d**, **Supplementary Table 2**).

To better understand the activation mechanisms with KHF waveforms, we investigated the effect of their duration on membrane potential changes and threshold currents in our computational model. Simulations of axonal membrane potentials indicate that the negative phases of the KHF waveforms actively counteracted the depolarization by the preceding positive phase that would have otherwise resulted in the generation of an action potential (**Supplementary Fig. 3**). To overcome this counteracting effect, either the stimulation current or the number of stimulation pulses had to be increased. Consistent with our experimental results, we observed that afferent and efferent fibers required less current with increasing KHF waveform duration to evoke responses at threshold (**Fig. 2e**). Moreover, we found that response latencies increased with increasing KHF waveform duration (**Fig. 2f**).

### KHF waveforms increase the co-activation of afferent and efferent fibers

Neuromodulation of spinal circuits by tSCS requires the recruitment of afferent fibers to enable the interaction with spinal circuits^20^, which is thought to be necessary for promoting neuroplasticity and functional recovery^23–25^. To examine the relative recruitment of afferent vs. efferent fibers, we used threshold stimulation currents and maximal responses (**Fig. 3a,b**). We quantified the ratio of stimulation currents required to evoke H-reflexes vs. M-waves at their respective thresholds, where an M/H stimulation current ratio larger than 1 would indicate that H-reflex responses are elicited at lower thresholds than M-waves, and a value smaller than 1 would indicate that M-waves are recruited first. We found that waveform had a significant effect on the M/H threshold ratio, with larger ratios found for the conventional waveforms than for each of the KHF waveforms (**Fig. 3a**, **Supplementary Table 1**). This indicates that at threshold currents, conventional waveforms recruit a larger proportion of afferent fibers than motor fibers compared to KHF waveforms. Importantly, there was no significant difference between KHF waveforms of different durations, suggesting that an increased duration did not change the relative afferent-to-efferent recruitment over a single 100 µs biphasic pulse. Notably, the conventional pulse of 2 ms duration was the only waveform for which the ratio was significantly larger than 1 (mean ratio 1.305, 95%-CI: 1.13 to 1.48), while all ratios with KHF waveforms were significantly lower than 1. All KHF waveforms result in the elicitation of a non-synaptically transmitted M-wave at lower currents than the H-reflex.

**Figure 3.**
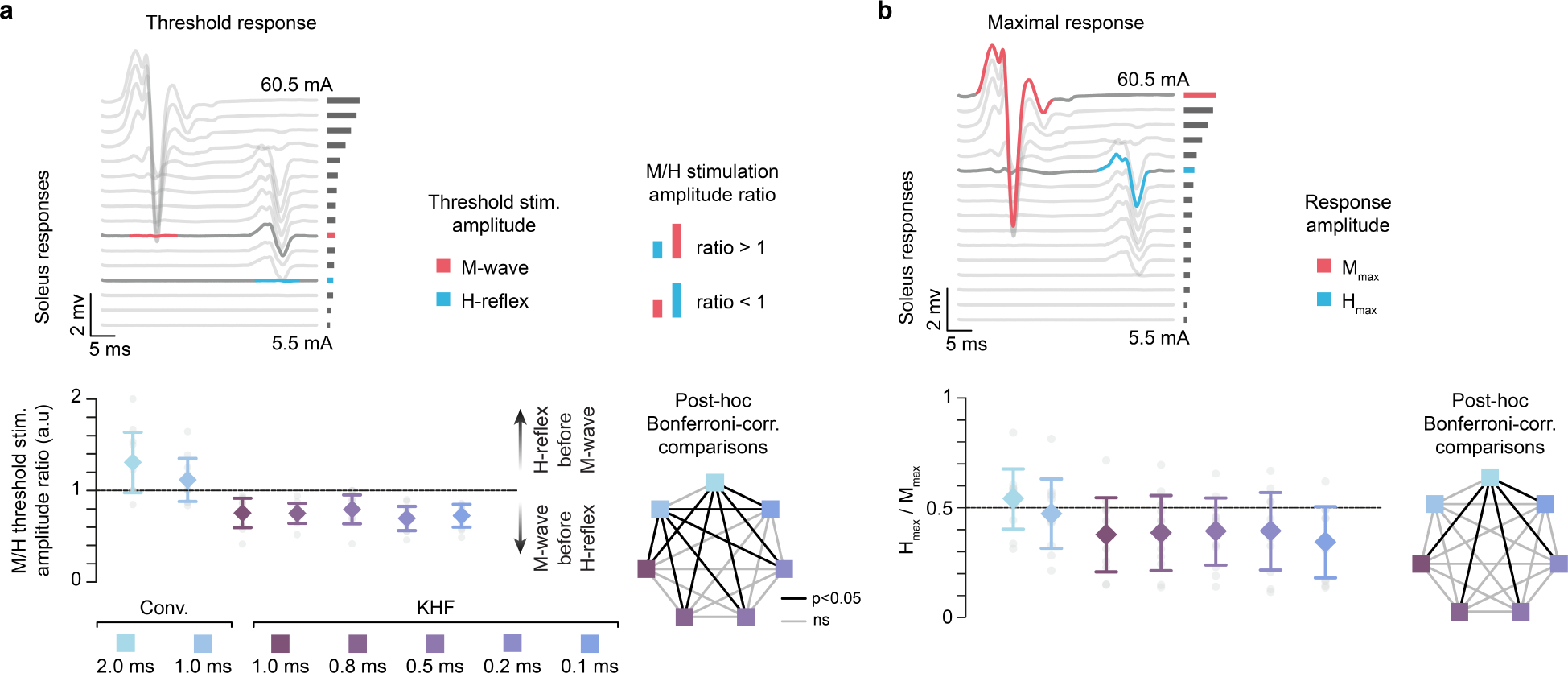
KHF waveforms preferentially recruit motor fibers. **a**. We compared threshold stimulation currents to evoke responses in the soleus muscle for conventional and KHF waveforms. We used the M/H stimulation threshold ratio as a measure of the relative recruitment of efferent to afferent fibers. A ratio greater than 1 indicates that the H-reflex occurs at lower stimulation currents than the M-wave. **b.** We used the H_max_/M_max_ ratio as a measure of the relative recruitment of afferent and efferent fibers. Lower H_max_/M_max_ values indicate a lower proportion of reflex-activated motor neurons in the soleus. Central diamonds represent the estimated means while whiskers indicate the 95% confidence intervals. The heptagon indicates the results for the post-hoc Bonferroni-corrected comparisons between waveforms; the solid black line indicates statistically significant differences between waveforms (p < 0.05). Abbreviations: stimulation (stim.), corrected (corr.), arbitrary units (a.u.), kilohertz frequency (KHF), not significant (n.s.). Evoked responses shown from 2 ms conventional waveforms are shown for RS001.

The increased co-activation of efferent fibers relative to afferent fibers with KHF waveforms would increase the amount of antidromic collisions with reflex-mediated responses. We compared the ratio between the maximum H-reflex and M-wave amplitudes that could be generated by each type of waveform with the H_max_/M_max_ ratio as a measure of the highest proportion of the soleus motor neuron pool that can be reflex-activated. We would expect that the increase in antidromic collisions would dampen the maximum amplitudes achievable by H-reflexes, resulting in a reduced H_max_/M_max_ ratio. Waveform had a significant effect on the H_max_/M_max_ ratio (**Fig. 3b**, **Supplementary Table 3**), with the conventional 2 ms waveform having a larger ratio than any of the KHF waveforms, and no KHF waveforms being significantly different from one another. Waveform had a significant effect on H_max_, but not on M_max_. However, none of the post-hoc comparisons were significant (**Supplementary Table 3**).

Electrophysiological recordings of H-reflexes and M-waves offer a way to compare the relative recruitment of afferent and efferent fibers. However, they do not provide absolute counts of the recruitment of these fibers. This is because a single recruited motor fiber can trigger an M-wave, whereas many recruited afferent fibers are required to generate an H-reflex^57,58^ (**Fig. 4a**). We used our computational model to understand the appearance of M-waves at lower stimulation currents with respect to H-reflex responses in KHF waveforms. We found that although both conventional and KHF waveforms preferentially recruit proprioceptive afferents (**Fig. 4b**), there was a larger overlap in the simultaneous recruitment of efferents and proprioceptive afferents with KHF waveforms (**Fig. 4c**). In KHF waveforms, the recruitment of efferent fibers started to occur at stimulation intensities in which the occurrence of H-reflexes may be triggered by afferent recruitment. Here, we refer to the stimulation window in which afferent recruitment will likely result in an H-reflex as the H-threshold probability window (> 20% recruited afferent fibers). Our model showed no recruitment of motor fibers within the H-reflex probability window for conventional waveforms, but a non-zero recruitment of efferent fibers within the window for KHF waveforms (**Fig. 4c**). Therefore, the probability of direct motor responses for KHF waveforms becomes more likely than trans-synaptic activation of motor neurons through proprioceptive afferents alone (**Fig. 4d,e**).

**Figure 4.**
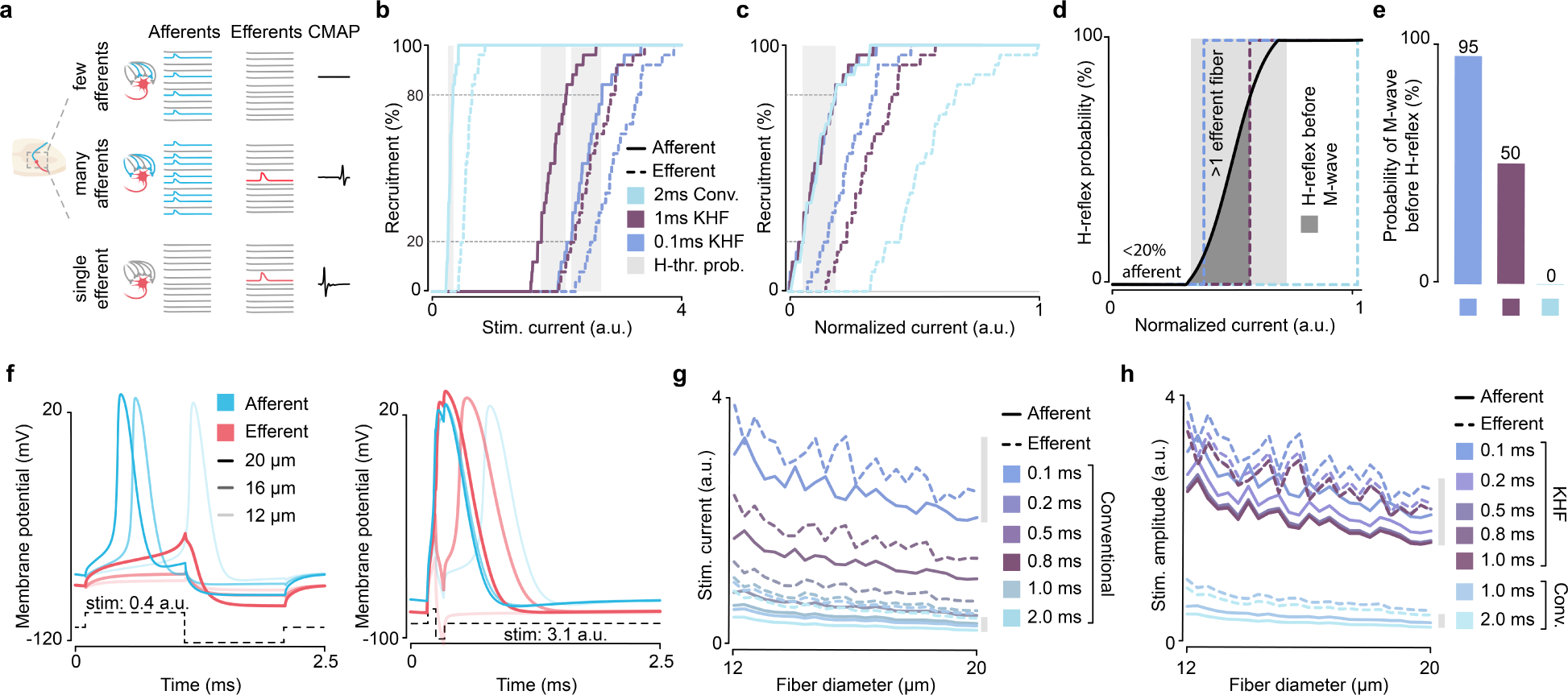
Recruitment of efferent fibers during the H-reflex window increases the likelihood of M-wave at threshold intensities with KHF waveforms. **a**. Recruitment of afferent and efferent fibers in a 25-to-1 afferent-to-efferent computational synaptic model. For this model, more than 8 afferent fibers had to be activated to elicit an action potential in the efferent fiber. Inactivated efferent fibers and CMAP responses included for illustration purposes, as these were not part of the model. **b**. Recruitment of efferent and afferent fibers as a function of stimulation current. Note that the thresholds and H-threshold probability windows vary across waveforms. **c**. Recruitment of efferent and afferent fibers with currents normalized to have matching H-threshold probability windows across waveforms. Even though the probability windows and the recruitment of afferent fibers overlap, the relative recruitment of efferent fibers varies between waveforms. Note that in the 2 ms conventional waveform, there is no efferent recruitment within the H-threshold probability window **d**. Probability of eliciting an H-reflex for each waveform. The dark gray area represents the currents at which there is a probability that the H-reflex will be elicited before the M-wave. The light gray area represents the H-threshold probability window. **e**. Probability of eliciting an M-wave before an H-reflex across waveforms. **f.** Membrane potentials in fibers of different diameters during stimulation with 2 ms and 0.1 ms conventional waveforms. Long-duration waveforms allow for recruitment of afferent fibers without recruitment of efferent fibers. **g**. Recruitment of fibers with different diameters across stimulation currents for conventional waveforms. Gray vertical bars indicate the range in currents for recruitment of proprioceptive fibers with different diameters for the 0.1 ms and 2 ms waveforms. Note that with conventional 2 ms pulses, proprioceptive afferents of different diameters can be recruited at a wide range of stimulation currents without also recruiting motor fibers. This is not the case with the 0.1 ms pulse. **h**. Recruitment of fibers with different diameters across stimulation currents for waveforms that were tested in the peripheral nerve stimulation experiments. Gray vertical bars indicate the range in currents for recruitment of proprioceptive fibers across stimulation currents in 1 ms KHF and 2 ms conventional waveforms. Abbreviations: compound muscle activation (CMAP), kilohertz-frequency (KHF), conventional (conv.), stimulation (stim.), arbitrary units (a.u.), H-threshold probability window (H-thr. prob.).

### Long pulse durations allow for the recruitment of a larger proportion of proprioceptive fibers without the simultaneous recruitment of motor fibers

To gain a deeper understanding of how pulse duration changes the shape of the recruitment curves for afferent and efferent fibers, we used our computational model to compare the extracellular membrane potentials generated by stimulation with conventional pulses of 2 ms and 0.1 ms duration in fibers of different diameters (**Fig. 1b**, tibial nerve). To accurately simulate the biophysical differences between afferent and efferent fibers, we used two distinct axon models that incorporate their physiological differences as a function of fiber diameter^56^ (**Supplementary Fig. 4**).

We found that the long 2 ms waveform allows for the slow depolarization of afferent fibers with different diameters, resulting in the recruitment of a larger proportion of afferent fibers without depolarization of motor fibers (**Fig. 4f**). In turn, the recruitment of the same proportion of afferent fibers with different diameters with a short 0.1 ms conventional waveform requires higher stimulation currents (**Fig. 4f**). The faster depolarization results in the simultaneous recruitment of efferent fibers with large diameters (**Fig. 4f**, **Supplementary Fig. 4c**). While there is a wide range of currents for conventional 2 ms waveforms where it is possible to recruit afferent fibers of different diameters without the recruitment of efferent fibers, there is only a short range of stimulation amplitudes where this is possible for the short 0.1 ms pulses (**Fig. 4g**). Similarly to shortening pulse duration, the use of KHF waveforms resulted in the simultaneous recruitment of afferent and efferent fibers across a wide range of stimulation currents (**Fig. 4h**). The difference in recruitment between conventional and KHF waveforms was much larger than those between KHF waveforms of different durations (**Fig. 4h**).

Notably, matching parameters in afferent fibers related to the hyperpolarization-activated cyclic nucleotide-gated (HCN) channels (conductance, rate constant, or all HCN-related parameters) to those in efferent fibers^56,59^ brought the resting membrane potentials closer together and largely reduced differences in their depolarization behavior (**Supplementary Fig. 4c**). This suggests that HCN-related parameters alone accounted for a large part of the differences in afferent vs. efferent depolarization by long or short duration pulses.

Results from the computational model suggest that although both conventional and KHF waveforms of different durations preferentially recruit proprioceptive afferents, KHF waveforms impact the relative activation of afferent and efferent fibers. The downstream effect of these differences in afferent vs. efferent recruitment results in a scenario where direct motor responses are more likely to occur with KHF waveforms, as reflected by the results from the peripheral nerve stimulation experiments.

Recruitment of afferent fibers by neurostimulation is an essential component for neuromodulation^24^. Studies in computational modeling and human neurophysiology have suggested that muscle responses elicited with conventional waveforms by eSCS and tSCS of the lumbar spinal cord are mediated by the preferential recruitment of proprioceptive fibers within the posterior roots^17,18^. However, the activation mechanisms of KHF waveforms in tSCS remain poorly understood. Therefore, we next sought to understand how the reduced efficacy of KHF waveforms in recruiting afferent fibers translates into the application of lumbar and cervical tSCS.

### Afferent recruitment in tSCS depends on the waveform and is compromised in cervical stimulation

In lumbar tSCS, the elicitation of posterior root-muscle (PRM) reflexes in lower limb muscles, i.e., spinal reflexes that share essential characteristics with the classical H-reflex^9,20^, has been used as an electrophysiological marker for local activation of lumbar proprioceptive afferents^17,60,61^. Paired-pulse paradigms are used to assess post-activation depression of evoked responses, which affect reflex-mediated responses but not direct muscle responses^1,20,21^. To determine the extent to which evoked responses in tSCS are mediated by the recruitment of proprioceptive afferent fibers, we applied paired pulses of the 2 ms conventional and the 1 ms KHF waveforms in both lumbar and cervical tSCS (**Fig. 5a, b**). We hypothesized that if responses were mediated by proprioceptive afferents, responses to the test (second) stimuli would show significant suppression due to post-activation depression (**Fig. 5c**). If, however, responses were mediated by direct recruitment of efferent fibers, the responses to the test stimuli would lack suppression.

**Figure 5.**
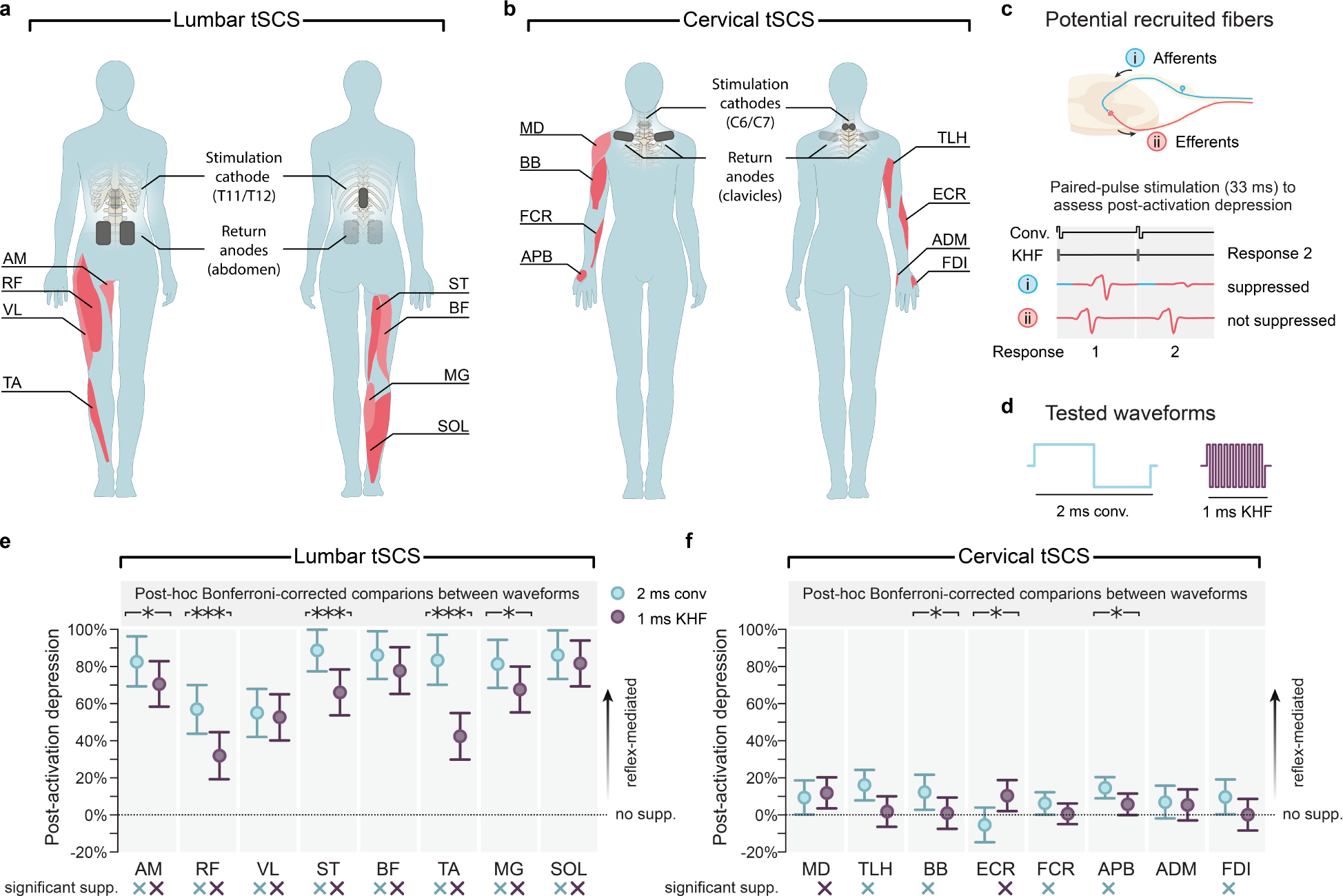
Suppression by paired-pulse stimulation reveals a low degree of post-activation depression with KHF waveforms and in cervical tSCS. **a**. Stimulation electrodes and EMG recordings for lumbar tSCS. **b**. Stimulation electrodes and EMG recordings for cervical tSCS. **c**. To understand the degree of afferent recruitment by conventional and KHF waveforms, we used a paired-pulse stimulation paradigm and evaluated the amount of post-activation depression by quantifying the suppression of the response to the second stimulation pulse. Reflex-mediated responses would result in a significant amount of suppression, whereas no suppression would be observed in the recruitment of efferent fibers bypassing the synapse. **d**. We evaluated the 2 ms conventional waveform and the 1 ms KHF waveform. **e**. Amount of post-activation depression across muscles in lumbar tSCS. **f.** Amount of post-activation depression across muscles in cervical tSCS. Central circles represent the estimated means while whiskers indicate the 95% confidence interval. The asterisks on the top of each muscle indicate the results for the post-hoc Bonferroni-corrected comparisons between waveforms; the × below the muscles indicate a significant amount of post-activation depression. ∗ p < 0.05; ∗∗∗ p < 0.0001. Abbreviations: Kilohertz frequency (KHF), adductor magnus (AM), rectus femoris (RF), biceps femoris (BF), semitendinosus (ST), vastus lateralis (VL), tibialis anterior (TA), medial gastrocnemius (MG), and soleus (SOL), medial deltoid (DM), biceps brachii (BB), triceps long head (TLH), flexor carpi radialis (FCR), extensor carpi radialis (ECR), abductor pollicis brevis (APB), abductor digiti minimi (ADM), and the first dorsal interosseous (FDI), suppression (supp.).

Suppression of the test responses in lumbar tSCS was significantly affected by the factors waveform and muscle (**Supplementary Table 4)**. Across muscles, amplitudes of the test responses were reduced by 77.6% (95%-CI: 88.4% to 66.8%) with the conventional waveform and by 61.3% (72.0% to 50.5%) with the KHF waveform. Importantly, suppression was significant for all muscles and both waveforms tested (**Fig. 5e**, **Supplementary Table 4**), suggesting that responses with both waveforms in lumbar tSCS are largely mediated by proprioceptive afferents. However, we found that the amount of suppression was significantly lower with KHF than with conventional waveforms for five of eight muscles (**Fig. 5e**), suggesting that there was a larger proportion of motor efferent fibers being recruited relative to proprioceptive afferent fibers.

The level of post-activation depression can depend on response size, with smaller responses being more susceptible to suppression^62^. While both factors waveform and muscle were found to have had a significant effect on the peak-to-peak amplitudes of the conditioning (first) responses, post-hoc comparisons showed significant differences only in the semitendinosus, with smaller responses evoked by KHF than conventional waveforms (**Supplementary Table 4**).

In cervical tSCS, suppression of the test responses was significantly affected by the factor waveform, but not by the factor muscle (**Supplementary Table 4**). Across muscles, amplitudes of the test responses were reduced by 8.7% (95%-CI: 12.5% to 4.8%) with the conventional waveform and by 4.6% (8.3% to 0.9%) with the KHF waveform. Suppression was significant for five of eight muscles with the conventional waveform, and for two of the eight muscles with the KHF waveform (**Fig. 5f**, **Supplementary Table 4)**. Both factors waveform and muscle affected the size of the conditioning responses (**Supplementary Table 4**). Smaller responses to KHF than conventional waveforms were found in the case of the abductor digiti minimi, biceps brachii, and the first dorsal interosseous. Despite the statistically significant suppression in some muscles by cervical tSCS, the small degree of suppression suggests that recruitment of efferent fibers plays a substantial role in the generation of these responses.

Compared to lumbar tSCS, the amount of suppression due to post-activation depression across muscles was significantly lower in cervical tSCS (**Fig. 5e,f**), with both factors waveform and stimulation site being significant (**Supplementary Table 5**).

### Elicited responses are mediated by the recruitment of posterior roots in lumbar tSCS and anterior roots in cervical tSCS. KHF waveforms make recruitment of posterior roots less likely

As an additional measure to identify whether conventional and KHF waveforms in lumbar and cervical tSCS recruit afferent or efferent fibers, we compared latencies between responses elicited by tSCS and peripheral conduction times calculated from M-wave and F-wave responses to tibial nerve stimulation in the medial gastrocnemius (**Fig. 6a**), and to median nerve stimulation in the abductor pollicis brevis (**Fig. 6b**). Peripheral conduction time represents the time it takes an action potential to travel from the motoneuronal cell bodies to the muscle^63,64^. We hypothesized that recruitment of afferent fibers in tSCS would lead to response latencies longer than peripheral conduction time. In contrast, recruitment of efferent fibers in tSCS would lead to response latencies equal to or shorter than peripheral conduction time.

**Figure 6.**
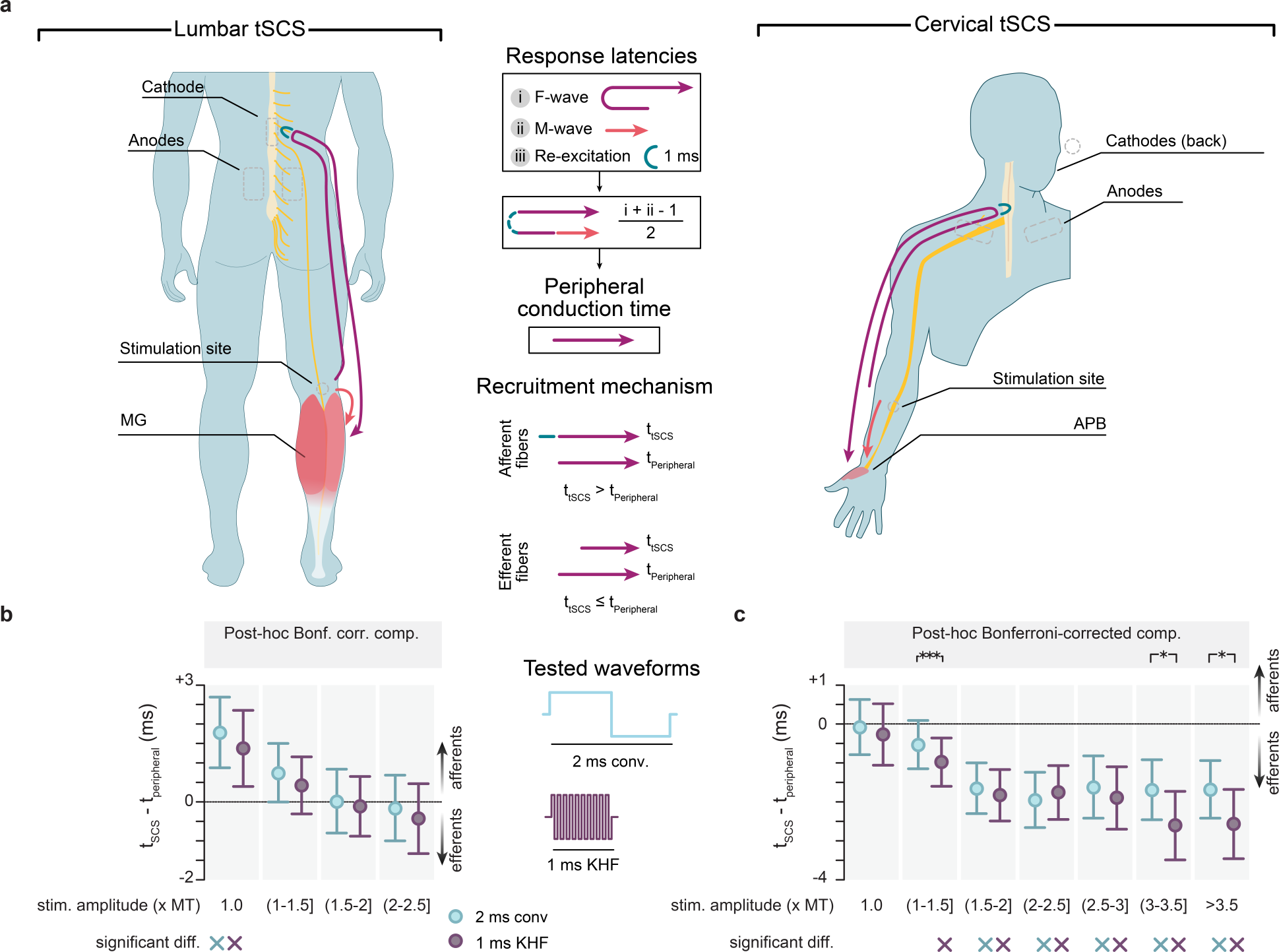
Comparisons of response latencies to peripheral conduction time reveal marked differences in recruitment mechanisms between lumbar and cervical tSCS. **a**. M-waves and F-waves were recorded at the stimulation amplitude to evoke a motor response at threshold and 1.25×M_max_, respectively. Peripheral conduction time was computed using the noted equation, where *i* represents the time that it takes for an F-wave to travel antidromically from the stimulation site to the anterior horns and orthodromically back to the muscle^66^, *ii* represents the time that it takes an M-wave to travel orthodromically from the stimulation site directly to the muscle, and 1 ms corresponds to the time required for an F-wave to re-excite the motor neuron soma^63,64^. Peripheral conduction time is an estimate of the time that it takes for an action potential to travel from the anterior horn of the spinal cord to the muscle^63,64^. **b**. Differences in response latency between tSCS and peripheral nerve stimulation are shown across stimulation currents for lumbar tSCS. Positive values indicate that responses evoked by tSCS had longer latencies compared to peripheral nerve stimulation and suggest that the recruitment mechanisms are mediated by posterior roots. **c**. Data for cervical tSCS. Negative values indicate recruitment at the anterior roots. Central circles in **b** and **c** represent the estimated means while whiskers indicate the 95% confidence interval. The asterisks on the top of each muscle indicate the results for the post-hoc Bonferroni-corrected comparisons between waveforms; the × below the muscles indicate that there is a statistically significant difference between t_tSCS_ and t_peripheral_. ∗ p < 0.05; ∗∗∗ p < 0.001. Abbreviations: medial gastrocnemius (MG), abductor pollicis brevis (APB), stimulation (stim.), muscle activation threshold (MT), difference (diff.), Bonferroni (Bonf.), corrected (corr.), comparison (comp.).

Stimulation current is an important factor that can influence the relative recruitment of afferent and efferent fibers (**Fig. 4**, **Supplementary Fig. 4**). However, while motor threshold is sometimes used to tune stimulation parameters ^7,61,65^, there is a lack of understanding of how stimulation amplitudes relative to motor threshold affect the relative recruitment of afferent and efferent fibers in tSCS. To investigate whether recruitment sites would change with increasing stimulation currents, we introduced different stimulation current bins: bin 1, containing responses evoked at threshold; bin 2, (1-1.5] x threshold; bin 3, (1.5-2.0] x threshold; bin 4, (2.0-2.5] x threshold; bin 5, (2.5-3.0] x threshold; bin 6, (3.0-3.5] x threshold; and bin 7, >3.5 x threshold.

In lumbar tSCS, we found a significant effect of stimulation current (considering bins 1-4, **Supplementary Table 6**), but not stimulation waveform. Response latencies with lumbar tSCS at threshold (i.e., bin 1) were longer than the peripheral conduction time for both conventional (mean difference 1.785 ms, 95%-CI: 0.875 ms to 2.695 ms) and KHF (mean difference 1.372 ms, 95%-CI: 0.396 ms to 2.348 ms) waveforms (**Fig. 6b**). Although response latencies at high stimulation currents (bins 2-4) were not significantly different from peripheral conduction time (**Supplementary Table 6**), post-activation depression was still considerable at these currents (**Supplementary Table 7**). This suggests that although some efferent recruitment is present, a large portion of the response is reflex-mediated.

In cervical tSCS, both stimulation current (considering bins 1-7) and waveform affected the difference between peripheral conduction time and response onsets (**Supplementary Table 6**). Response latencies with cervical tSCS at threshold (bin 1) showed no statistically significant difference to peripheral conduction time for both waveforms (**Fig. 6c**). Response latencies at higher stimulation currents (bins 3-7 in conventional and bins 2-7 in KHF) were significantly shorter than peripheral conduction time (**Supplementary Table 6**). Moreover, the amount of post-activation depression for the abductor pollicis brevis was absent or weak across stimulation currents for both waveforms (**Supplementary Table 7**).

Together, these results suggest that although both waveforms preferentially activate afferent fibers in lumbar tSCS (**Fig. 6b**), KHF waveforms demonstrate a significantly lower degree of post-activation than conventional waveforms (**Fig. 5e**). Unexpectedly, the advantage of conventional waveforms to recruit afferent fibers does not translate into cervical tSCS, where the lack of suppression for both conventional and KHF waveforms (**Fig. 5f**), combined with shorter response latencies than peripheral conduction times (**Fig. 6c**), strongly suggests that recruitment of efferent fibers plays a major role.

We combined the differences in conduction times with conduction velocities in posterior and anterior roots to estimate the average recruitment sites (i.e., action potential generation site) for lumbar and cervical tSCS (**Supplementary Fig. 5**). Medial gastrocnemius recruitment sites with lumbar tSCS at threshold stimulation amplitudes were estimated to be 41mm distal to the proximal end of the posterior roots with conventional waveforms and 13 mm with KHF waveforms. Abductor pollicis brevis recruitment sites with cervical tSCS at threshold stimulation amplitudes were estimated to be at the proximal end of the anterior roots with both waveforms. However, the recruitment sites for cervical tSCS moved distally along the efferent pathways with increasing stimulation pathways, with stimulation sites migrating from 5 to 14 cm towards the periphery.

### Kilohertz-frequency waveforms require higher stimulation intensities to elicit responses in tSCS

In agreement with our peripheral nerve stimulation experiments and previous studies^40,41^, we observed that KHF waveforms required higher current and charge than conventional waveforms in both lumbar and cervical tSCS (**Supplementary Table 8**). Specifically, KHF waveforms required on average 5.0 ± 0.6 times more current than conventional waveforms in lumbar tSCS, and 4.1 ± 0.4 times more current in cervical tSCS. Concurrently, KHF waveforms required on average 2.5 ± 0.3 times more charge than conventional waveforms in lumbar tSCS, and 2.0 ± 0.2 times more charge for cervical tSCS.

## Discussion

The original motivation for the development of tSCS was to non-invasively activate proprioceptive afferent fibers in the posterior roots to provide a neuromodulatory input to the lumbar spinal circuitry, similar to lumbar eSCS^16,36,67^. Computer simulations conducted in coupled volume conductor and neural axon models of the thoraco-lumbar spine indicated that most of the current induced by tSCS flows around the spine. However, a fraction of the current crosses the vertebral canal by primarily passing through ligaments and intervertebral discs, creating relatively large current densities within the cerebrospinal fluid where the roots are immersed^37,38,68^. Taken together with changes in the orientation of the posterior rootlets within the electric field and local inhomogeneities in electrical conductivity at their entry into the spinal cord, these computational investigations proposed that similar to eSCS, lumbar tSCS with conventional waveforms predominantly activates proprioceptive afferent fibers in the posterior roots^11,37,69^. Moreover, the use of long-duration stimulation waveforms was found to further prioritize recruitment of afferent fibers by maximizing the difference in the strength-duration properties between motor and proprioceptive axons^70^.

Importantly, these biophysical foundations of lumbar tSCS may not directly translate to alternative stimulation paradigms, such as KHF waveforms, or to the cervical spine^10,32,71^. These KHF waveforms were first introduced by Gerasimenko, Edgerton, and colleagues in tSCS, as 0.5 ms or 1.0 ms KHF waveforms with a carrier frequency of 5 or 10 kHz and amplitudes up to 200 mA^10,71^, with the rationale that such stimulation would not elicit pain even when applied at energies required to reach spinal networks^10^. However, KHF waveforms have been recently shown to require much higher stimulation current and charge to elicit muscle responses than conventional waveforms^40,41^. Therefore, potential pain relief benefits by KHF waveforms are completely abolished when controlling for the magnitude of the evoked response^40,41^.

From a biophysical point of view, using KHF waveforms to activate neural axons appears counterintuitive, as high-frequency alternating currents should de-, and subsequently re-, and hyperpolarize the membrane in quick succession with each AC cycle. Excitation caused by KHF waveforms was originally described to work through summation processes^72^. Specifically, the effect of the first positive half-wave was thought not to be completely canceled by the subsequent negative half-wave of an AC cycle. Consequently, after several AC cycles, the membrane potential was thought to be sequentially depolarized, eventually reaching the threshold for action potential generation, an effect initially introduced in 1944 as the Gildemeister effect^73^ and later described in detail in animal experiments^49^.

Building upon these investigations in animals 80 years ago, our work provides evidence that KHF waveforms applied by tSCS in humans activate fibers through summation processes. Specifically, the decreasing thresholds and the increasing onset latencies of H-reflexes with increasing KHF burst duration (**Fig. 2c,d**), substantiate the claim that proprioceptive afferent fibers are activated at increasing numbers of AC cycles. Furthermore, the 1 ms estimated onset latency difference between H-reflexes evoked by the 1.0 ms and 0.1 ms KHF waveforms suggests that the longer KHF waveform brings the proprioceptive fibers to firing threshold only at the last AC cycle within the burst. Our computational investigation explains this finding and shows that membrane depolarization by the positive phases is sequentially interrupted by the negative phases of each AC cycle, thus delaying action potential generation towards the last cycle at threshold (**Supplementary Fig. 3**).

The counteracting effects of sequential hyperpolarization may also explain the considerably higher current and charge required to elicit responses with KHF waveforms (**Fig. 2c** and **Supplementary Table 8**). The four to five times higher stimulation intensities for KHF waveforms to evoke muscle responses in our neurophysiological experiments and computational model (**Fig. 2**, **Supplementary Fig. 2**, and **Supplementary Table 1**) are in line with previous observations^40,41^. One study was unable to reach response threshold with KHF waveforms at current amplitudes up to 200 mA^74^, and another described biphasic KHF waveforms as ‘intentionally ineffective’ sham for lumbar tSCS, as such stimulation would not produce motor output in the lower limbs^75^. These observations combined with our reported results strongly suggest that a shift is necessary in the standard of reporting stimulation currents used for rehabilitation^7,76^, as current alone provides insufficient information without a respective muscle activation threshold for normalization. We argue that reporting muscle activation threshold in addition to stimulation current is necessary to correctly interpret stimulation intensities that can lead to improved recovery outcomes.

Our computer simulations showed that KHF waveforms recruit fewer proprioceptive fibers relative to motor fibers than conventional pulses (**Fig. 4d-h**). These computational observations were in line with our neurophysiological experiments, where we found that M/H threshold stimulation ratios were significantly higher for conventional waveforms than for any KHF waveform (**Fig. 3a**). These findings have potentially negative implications for the use of KHF waveforms in clinical studies for neurorehabilitation, as the direct activation of proprioceptive afferents is believed to be essential for functional recovery^24,25^.

The M/H ratios did not show significant differences between KHF waveforms. The studied KHF waveforms were composed of a series of 0.1 ms biphasic pulses, i.e., short conventional waveforms, which have long been recognized to result in higher thresholds and lower amplitudes in H-reflexes compared to longer conventional waveforms of 0.5-1 ms^44,45^. Our experimental and computational model results suggest that while long burst durations in KHF waveforms (1 ms), as currently used in clinical studies^7,33,77–79^, can lower the muscle response threshold over short burst durations (0.1 ms), lengthening KHF waveform duration does not offer a significant advantage in the relative recruitment of motor and proprioceptive fibers. Importantly, our computational model provides not only a simulated observation of discrepancies in proprioceptive vs. motor fiber recruitment with KHF and conventional waveforms but also a mechanistic framework to understand these differences. While summation processes do occur across each cycle of the KHF waveform, the depolarization of the axonal membrane can be interrupted by the negative phase of each AC cycle (**Supp. Fig. 3**), leaving single 50 µs depolarization phases as the dominating characteristic of the KHF waveforms (**Fig. 4g,h**). Due to intrinsic membrane properties such as differences in resting membrane potential^56^ (**Supplementary Fig. 4c**), motor fibers are more likely to be recruited simultaneously to proprioceptive fibers with short waveforms (**Fig. 4b–e**). This aligns with the strength-duration curves showing that shorter pulses recruit more motor fibers relative to proprioceptive fibers^43,44,48^. Although H-reflex experiments and computational modeling allowed us to understand the mechanisms underlying the recruitment of motor and proprioceptive fibers by conventional and KHF waveforms, they do not fully reveal the clinical potential for their use in lumbar and cervical tSCS. Anatomical differences between the cervical and lumbar spines—such as the projection and overlap of cervical spinous processes, thinner intervertebral discs in the cervical region, and distinct orientations and curvatures of spinal roots—affect current flow directions and the distribution of induced currents^37,38,69,80,81^. These anatomical variations likely result in different locations of low-threshold hotspots and varied excitation of proprioceptive and motor fibers.

In lumbar tSCS, suppression of the test response in paired-pulse paradigms has been used as an electrophysiological marker for local activation of lumbar proprioceptive afferents ^17,61,80^. While a response suppression of 77.6% by paired conventional waveforms in lumbar tSCS in our study is compatible with earlier observations^36,68,82^, the suppression of 61.3% by KHF waveforms suggests an increased proportion of efferent motor fiber recruitment.

Intriguingly, in cervical tSCS, while the conventional waveforms still produced greater suppression of the second responses than the KHF waveforms (8.7% vs. 4.6% reduction), the weak or even completely absent post-activation depression in most muscles demonstrated significantly weaker suppression for cervical than lumbar tSCS with both waveforms (**Fig. 5**). Our results corroborate previous observations that conventional waveforms in cervical tSCS with cathodes placed over the cervical spinous process and various anode configurations (one anode over the spine placed caudally to the cathode, two anodes placed bilaterally either over the clavicles or the iliac crests) did not result in a statistically significant suppression of the second responses in the upper extremity muscles^60^. Only a configuration with an anode placed over the anterior neck applying conventional pulses demonstrated suppression of the second response, although suppression was still incomplete and roughly amounted to 50% of the first responses^60,83,84^. However, delivering high-intensity stimulation continuously over anterior neck muscles during rehabilitation may pose significant feasibility challenges. It is important to note that current applications in tSCS vary widely in the placement for stimulating electrodes^7,32,60,79,85–88^. Recruitment mechanisms should be carefully studied in the proposal of novel electrode placements.

Theoretically, the absence of post-activation depression in cervical tSCS could also be explained by assuming that upper limb PRM reflexes are less affected by presynaptic inhibition of Ia afferent terminals and by reduced neurotransmitter release upon repeated activation^60,83^. However, at interstimulus intervals of 60 ms or less, comparable to the interstimulus intervals used in paired tSCS studies, the second flexor carpi radialis H-reflex was shown to be suppressed by 90% of the first reflex^89^, compared to the suppression by only 6% when evoked by cervical tSCS using conventional waveforms in this study. Hence, differences in the excitability between cervical and lumbar reflex circuits are an unlikely explanation for the incomplete or absent post-activation depression in cervical tSCS, and biophysical differences may rather explain this discrepancy.

The longer onset latencies of medial gastrocnemius responses by tSCS compared to peripheral conduction time (1.8 ms for conventional and 1.4 ms for KHF waveforms, **Fig. 6b**), further affirms that lumbar tSCS recruits afferent fibers within the posterior roots at threshold. In contrast, the absence of significant difference between latencies of abductor pollicis brevis responses by tSCS and peripheral conduction time suggest that cervical tSCS recruits motor fibers within the most proximal portions of the anterior roots. The further reduction in response latency at higher stimulation currents in cervical tSCS (up to 2.0 ms for conventional and 2.6 ms for KHF waveforms, **Fig. 6c**), which has been previously observed^33^, suggests that stimulation sites migrate peripherally toward the brachial plexus with increasing stimulation intensities. Although identifying the precise stimulation sites was outside the scope of this study, computational modeling can help in this endeavor.

## Conclusion

KHF waveforms were originally developed to maximize limb torques through neuromuscular stimulation, i.e., motor axon stimulation, at tolerable perception levels^90,91^. When KHF waveforms were adopted for tSCS, the idea presented was to neuromodulate spinal circuits painlessly^10,71^. The notion that KHF waveforms, at functional levels of stimulation currents, induce less pain than conventional waveforms has been long challenged in neuromuscular stimulation^92,93^ and more recently in tSCS^40,41,74^. Here, we revealed major biophysical drawbacks of KHF waveforms in the recruitment of proprioceptive fibers by neurophysiological studies in peripheral nerve stimulation, cervical and lumbar tSCS, and computational modeling. We showed that the disadvantages of KHF waveforms translate to lumbar tSCS, but surprisingly, in cervical tSCS, both conventional and KHF waveforms predominantly evoked motor responses non-synaptically. Our work challenges the continued promotion of KHF waveforms for tSCS and reveals limitations of current cervical tSCS applications. We thus posit that tSCS paradigms that aim to restore motor function should be underscored by a solid neurophysiological understanding of their underlying mechanisms of action before clinical translation.

## Methods

### Participants

This study was reviewed and approved by the Institutional Review Board of Washington University in St. Louis. Twenty-five unimpaired participants provided their informed consent to participate in this study, with 15 of those participating in the peripheral nerve stimulation experiment. However, three participants were excluded from the analysis due to the use of a monophasic, rather than biphasic, waveform, for a total of 12 included participants (8 female, 3 male, 1 preferred not to identify, average age 25.58 ± 3.52 years old). 10 in the cervical tSCS experiment (5 female, 4 male, 1 preferred not to identify, average age 24.3 ± 1.62 years old), and 10 in the lumbar tSCS experiment (5 female, 5 male, average age 26.7 ± 3.58 years old). Participant demographics can be found in **Supplementary Table 9**.

### Peripheral Nerve Stimulation Experiments

#### Experimental setup

We evaluated responses to tibial nerve stimulation by different stimulation waveforms in the right leg. We placed a round 3.2 cm diameter stimulation cathode at the popliteal fossa (all stimulating electrodes were PALS Neurostimulation Electrodes, Axelgaard Manufacturing Co., Ltd, USA), and a 5 x 9 cm anode at the lower patella. EMG responses were recorded from the soleus using silver chloride wet electrodes (all recording electrodes are Norotrode 20 bipolar, silver-silver chloride, disposable, SEMG electrodes), placed according to the SENIAM convention^94^. Prior to electrode placement, the skin was prepared with abrasive gel (NuPrep^®^, Weaver and Co., USA), applied with a Q-tip^®^ in circular motions, and wiped clean with an alcohol pad. Participants were then seated in a Biodex System 4 Pro^TM^ isokinetic dynamometer (Biodex Medical Systems, Shirley, NY, USA) with the right hip, knee, and ankle joints at approximately 120°, 110°, and 90°, respectively.

#### Data acquisition

Muscle responses were recorded using a wired connection to a Digitimer D360 amplifier (Digitimer Ltd, UK) using a 10-5000 Hz analog bandpass filter and a gain of 500. The responses were then fed through a Hum Bug noise eliminator (Hum Bug, Quest Scientific, India) to reduce 50-60 Hz line noise^95^. The filtered signals were digitized at 50 kHz using an analog-to-digital converter (Qualysis Analog Interface 16 Channels, Qualisys, Sweden) and then sent to a host computer running Qualisys Track Manager (v2021.1, build 6350). Custom software was developed in-house (Python 3.10.4) with the Qualisys Python SDK (v2.1.1) to communicate with the Qualisys Track Manager and record and visualize the data in real-time.

#### Experimental conditions

We tested 7 stimulation waveforms in peripheral nerve stimulation (**cf. Fig. 1**). All waveforms were biphasic: i) conventional, with a positive phase of 1 ms followed by a negative phase of 1 ms, referred to as a 2 ms conventional waveform, ii) KHF waveform with a 1 ms pulse modulated by a 10 kHz carrier wave (1 ms KHF), iii) conventional waveform with a positive phase of 0.5 ms and a negative phase of 0.5 ms (1 ms conventional) to account for differences in charge between the conventional and KHF waveforms, iv-vii) KHF waveforms with burst durations of 0.8 ms, 0.5 ms, 0.2 ms, and 0.1 ms. Note that the 0.1 ms KHF waveform at 10 kHz is equivalent to a 0.1 ms conventional waveform.

Stimulation was delivered using a constant current stimulator (Digitimer DS8R, Digitimer Ltd, UK) with modified firmware (D128R-201-IC1-RUSS-1.4.5.0) to enable the delivery of KHF waveforms. A National Instruments DAQ (USB-6001, National Instruments, USA) was used to send a trigger pulse to a Raspberry Pi Pico microcontroller (Raspberry Pi Pico, Raspberry Pi Foundation, UK) which would in turn send a trigger to the stimulator (single trigger pulses in conventional waveforms, or multiple trigger pulses 100 µs apart in KHF waveforms) to generate the pre-programed biphasic waveform. The DAQ was also used to digitally set the stimulation current for each waveform. The Pico’s ability to generate the proper waveform shapes was verified using a Keysight EDUX1052A digital storage oscilloscope connected to the DS8R stimulation leads.

#### Peripheral stimulation protocol

For each waveform, stimulation was first manually delivered by the experimenter to determine the stimulation currents in the soleus to evoke: i) H-reflexes at threshold, i.e., minimum stimulation currents to evoke H-reflexes with peak-to-peak amplitudes ≥ 50 µV, ii) M-waves at threshold, iii) H-reflexes with maximum attainable peak-to-peak amplitudes (H_max_), and iv) M-waves with maximum attainable peak-to-peak amplitudes (M_max_). The stimulation currents then used in the peripheral nerve stimulation protocol were set to correspond to 50% and 100% of the H-reflex threshold as well as to 5%, 25%, 50%, 75%, 100%, and 110% of the current required to evoke H_max_ for each waveform. This was repeated for the M-wave threshold and M_max_.

Three repetitions were performed at each stimulation current, for a total of 48 stimulation pulses per waveform. Pulses were delivered from lowest to highest stimulation current with 7±0.5 seconds (randomized) between pulses to avoid response suppression due to post-activation depression^17^. The testing order for the different waveforms was randomized at the start of each experiment (randarray, Microsoft Excel).

#### Data analysis and statistics

Data analysis was performed offline in MATLAB (Matlab R2020a, The MathWorks, Inc., Natick, MA) and IBM SPSS Statistics 28.0.1.1 for Windows (IBM Corporation, Armonk, NY). Statistical comparisons between waveforms were performed by fitting generalized linear mixed models (GLMMs) with subject as a random factor. α-errors of p < 0.05 (two-sided) were considered significant. Effect sizes are reported by the partial eta squared (*η*_*p*_^2^). All post-hoc tests were Bonferroni-corrected to adjust for multiple comparisons. Descriptive statistics are reported as mean ± SD or as estimated means along with SE or 95%-CIs.

EMG data was segmented using the stimulation trigger for each stimulation pulse (**Supplementary Fig. 1**). H-reflex and M-wave peak-to-peak amplitudes were computed as the difference between the maximum and minimum values of the EMG signal within the time windows of expected H-reflexes and M-waves. Mean values were computed per stimulation current. For each waveform, H-reflex and M-wave threshold stimulation currents were determined as the lowest currents that evoked H-reflexes and M-waves with peak-to-peak amplitudes ≥ 50 µV (**Fig. 2a**, and **Fig. 3a**). We then used these thresholds to compute the M/H threshold stimulation current ratio as a metric to compare the relative recruitment of motor and proprioceptive fibers by each waveform (**Fig. 3a**). Charges (µC) at H-reflex and M-wave thresholds were calculated by multiplying the stimulation current (mA) by the total duration of the positive phases of a given waveform, i.e., by 1 ms for the 2 ms conventional and by 0.5 ms for the 1 ms conventional waveforms and by 0.5 ms, 0.4 ms, 0.25 ms, 0.1 ms, and 0.05 ms respectively for the 1 ms, 0.8 ms, 0.5 ms, 0.2 ms and 0.1 ms KHF waveforms.

Peak-to-peak amplitudes of H_max_ and M_max_ (**Fig. 3b**) were used to compute the H_max_/M_max_ ratio for each waveform. For the determination of H-reflex latencies at threshold, mean slopes ± SD of the EMG were first calculated for 10 ms time windows preceding their expected onsets. H-reflex latencies were then calculated as the times between stimulus application (i.e., onset of the first positive phase of each waveform) and the first changes in EMG slope that exceeded the mean slopes by two times the respective SD (**Fig. 2d**).

### Computational model

All simulations were performed using the computational life-sciences simulation platform Sim4Life (version 7.2.1.11125, Zurich MedTech AG, Switzerland)^96,97^.

#### Volume conductor model

We generated a digital replica for the peripheral stimulation protocol based on the high-resolution Jeduk model of the Virtual Population Version 4.0 (IT’IS Foundation Virtual Population Version 4.0)^52–55^. For this purpose, we first manually regenerated the entire volume of the leg, filling holes, and connecting previously unconnected volumes. All tissues, except neural tissue, were assigned isotropic conductivity with dielectric properties sourced from the IT’IS Foundation low-frequency database. Neural tissues were assigned anisotropic conductivity, calculated as per previously established methods^6^. The volume conductor was discretized using structured modeling methods available in Sim4Life into 226.07 million voxels. Electrodes were manually placed on the skin analogous to the peripheral stimulation protocol. Electromagnetic simulations were performed using the Ohmic quasi-static low-frequency solver. Electrode boundary conditions were set to Dirichlet conditions, while the simulation domain boundaries were set to von Neumann conditions.

#### Neuron Simulation

Neural dynamics were simulated with the NEURON 7.8 solver (Yale University, Connecticut, USA) intrinsic to Sim4Life. The model included 75 motor and 75 sensory fibers, using the Motor MRG and Sensory MRG built into Sim4Life^56^. In the absence of histological information of afferent and efferent fiber diameter, all fiber diameters were initialized following a log-normal distribution (μ = 16.5 µm, σ = 2 µm). Where specified in the text, the fiber models were adjusted based on the sensory MRG model^56^, with adjustments made to channel conductance, rate constants, and Hyperpolarization-activated cyclic nucleotide-gated (HCN) channels for the Inter Node (STIN, MYSA, FLUT) regions using data from the Motor MRG model. Simulated waveforms replicated experimental conditions. The membrane potential was measured using the built-in Point sensor in Sim4life, which was manually positioned at the first activated node.

#### Recruitment curves

The stimulus amplitude is represented by the titration factor, which is a multiplier applied to the extracellular potential derived from electromagnetic simulation. The titration factor for each nerve fiber was determined using the least squares method and was defined as the minimum stimulus amplitude needed to trigger an action potential, where the change in the titration factor is less than 1%. The recruitment curve illustrates the percentage of nerve fibers activated at varying stimulus amplitudes.

#### H-threshold probability window

We digitally replicated the electrophysiological measurements of our peripheral stimulation protocol, focusing on the H-reflex and M-wave. First, we assumed that each directly stimulated motor efferent would produce a measurable M-wave. Second, we hypothesized that eliciting an H-reflex would require the recruitment of multiple proprioceptive afferents. Lacking a validated computational model for replicating the H-reflex with our neural dynamics simulations, we conservatively estimated the recruitment of proprioceptive fibers to be between 20% and 80%. We termed this range the H-threshold probability window (**Fig. 4b, c, d**). Specifically, when the recruitment percentage is below 20%, the H-reflex is assumed not to occur, whereas when it exceeds 80%, the H-reflex is assumed to have occurred. We normalized all H-threshold probability windows to overlap (**Fig. 4c** horizontal axis scaling factors: 2ms Conv.: 1, 1ms KHF: 0.22, 0.1ms KHF: 0.18). Within this probability window, we utilized a sigmoid curve^98^ (**Fig. 4d**), where the minimum probability corresponds to sigmoid(x=-2.5) and the maximum probability corresponds to sigmoid(x=2.5). The sigmoid function’s zero point aligns with the probability window’s center. We postulate that the M-wave is present once single motor fibers are recruited. Thus, the presence of the M-wave corresponds to a probability of 1 (**Fig. 4d**). Conversely, when the M-wave probability is 0, and the H-reflex probability is non-zero (**Fig. 4d**), it signifies that the H-reflex precedes the M-wave. When the M-wave probability is 1, and the H-reflex probability is less than 1, it indicates that the M-wave occurs before the H-reflex (**Fig. 4e**).

### Lumbar tSCS experiments

#### Experimental setup

We evaluated responses to lumbar tSCS by 2 ms conventional and 1 ms KHF waveforms in the right leg. We placed a rectangular 5 x 9 cm cathode centered on the T11 and T12 vertebrae, and two rectangular, interconnected 7.5 x 10 cm anodes on both sides of the navel. Vertebrae were identified via palpation and anatomical landmarks as described previously^8^. EMG responses were recorded from 8 right leg muscles: adductor magnus, rectus femoris, biceps femoris, semitendinosus, vastus lateralis, tibialis anterior, medial gastrocnemius, and soleus. Electrodes for the adductor magnus were placed by palpating for the muscle body as participants pulled their leg medially^99^, all other electrodes were placed according to the SENIAM conventions (no SENIAM guidance for adductor magnus)^94^. Skin preparation was the same as described for the peripheral nerve stimulation protocol. During the experiments, participants remained supine on a patient bed, with a small pillow placed under the head and neck to keep them comfortable. The participants’ arms were left relaxed and outstretched alongside their body. The participants’ legs were also relaxed and fully outstretched.

#### Data acquisition

Muscle responses were recorded using a wired connection to a Digitimer D360 amplifier with a 10-5000 Hz analog bandpass filter and a gain of 500. The responses of the rectus femoris and the medial gastrocnemius were fed through a HumBug noise eliminator to remove any 50-60 hz line noise^95^. The remaining channels did not have additional filtering. Signals were digitized at 25 kHz using a Qualysis branded digitizer and sent to a host computer running Qualisys Track Manager. The same in-house Python software as in peripheral nerve stimulation was used to record and visualize the data.

#### Experimental conditions

We tested 2 stimulation waveforms: i) the 2 ms conventional waveform, and ii) the 1 ms KHF waveform. Stimulation was applied using the same hardware as for the peripheral nerve stimulation protocol, set to deliver paired pulses with an interstimulus interval of 33 ms. The paired pulses were used to assess suppression of the responses to the second pulses and quantify the amount of post-activation depression^36,83^.

#### Lumbar tSCS protocol

For each waveform, stimulation was first manually delivered by the experimenter to determine the stimulation currents that evoke i) responses in the rectus femoris and medial gastrocnemius at threshold, i.e., minimum stimulation currents to evoke responses with peak-to-peak amplitudes ≥ 50 µV, and ii) rectus femoris and medial gastrocnemius responses with maximum attainable peak-to-peak amplitudes. The stimulation currents then used in the lumbar tSCS protocol were set to correspond to 50% and 100% of the current required to evoke responses at threshold as well as 5%, 25%, 50%, 75%, 100%, and 110% of the current required to evoke maximum responses in the two muscles.

Three paired pulses were applied at each stimulation amplitude, for a total of 48 repetitions per waveform. Pulses were delivered from lowest to highest stimulation current with 7±0.5 seconds (randomized) between repetitions. The testing order for the different waveforms was randomized (randarray, Microsoft Excel).

#### Data analysis and statistics

Data were analyzed using the same software, and statistical analyses were performed analogously to the peripheral stimulation protocol (GLMM models, alpha levels, post-hoc tests, corrections for multiple comparisons).

EMG data was segmented using the stimulation trigger of the first pulse of the paired-pulse stimuli. Peak-to-peak amplitudes for each of the two responses evoked by paired-pulse lumbar tSCS were calculated as the difference between the maximum and minimum values of the EMG signal within time windows of the first and the second pulses, respectively. Suppression due to post-activation depression was quantified by dividing the peak-to-peak amplitude of the second response by that of the first response, subtracting this difference from 1, and multiplying by 100%. A positive value would indicate suppression and a negative value would indicate facilitation.

For each waveform, threshold stimulation currents were determined for each muscle as the lowest currents that evoked responses to the first pulses of the paired stimuli with peak-to-peak amplitudes ≥ 50 µV. Charges at threshold were calculated by multiplying the stimulation current by the total duration of the positive phases of a waveform, i.e., by 1 ms for the 2 ms conventional waveform and 0.5 ms for the 1 ms KHF waveform. Response latencies were determined using the same analysis pipeline as for the peripheral nerve stimulation protocol, with time windows of 5 ms for the thigh muscles and 10 ms for the lower leg muscles to calculate the mean EMG slope ± SD prior to the expected response onset.

#### Peripheral conduction

Onset latencies of the responses in the medial gastrocnemius were compared to the peripheral conduction time of this muscle. Peripheral conduction time was determined by applying 1 ms biphasic pulses through a 3.2 cm diameter cathode placed over the tibial nerve in the popliteal fossa and a 3.2 cm diameter anode 2 cm proximal to evoke M-waves and F-waves in medial gastrocnemius. Stimulation was incrementally increased from the M-wave threshold. At 125% of M_max_, 60 pulses were applied every 3 ± 0.5 seconds (randomized) to evoke F-waves^100–102^. Peripheral conduction time of the medial gastrocnemius was then calculated as:

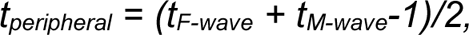

where *t_F-wave_* and *t_M-wave_* represent the F-wave and threshold M-wave latencies, respectively, and 1 ms is associated with the time for the orthodromically-traveling F-wave to re-excite the motoneuron at the soma^100^.

### Cervical tSCS experiments

#### Experimental setup

We evaluated responses to cervical tSCS by 2 ms conventional and 1 ms KHF waveforms in the right arm. We placed two round 3.2 cm diameter cathodes horizontally between the C6 and C7 vertebrae to target the motor neuron pools innervating the abductor pollicis brevis^9^, and two interconnected rectangular 5 x 9 cm anodes on the clavicles. The C6 and C7 vertebrae were identified using the flexion-extension palpation method^103^. EMG responses were recorded from 8 right-arm muscles: medial deltoid, biceps brachii, triceps long head, flexor carpi radialis, extensor carpi radialis, abductor pollicis brevis, abductor digiti minimi, and the first dorsal interosseous. The electrodes for the medial deltoid, biceps brachii, and triceps long head were placed according to the SENIAM conventions^94^. The remaining electrodes were placed according to the Atlas of Muscle Innervation Zones^104^ as SENIAM had no guidance for those muscles. Skin preparation was the same as for lumbar tSCS. During the experiment, participants remained supine on a patient bed, with a small pillow placed under the head and neck to keep them comfortable. The participants’ arms were left relaxed and outstretched alongside their body.

#### Data acquisition

Muscle responses were recorded using the same setup as for lumbar tSCS. The responses of abductor pollicis brevis and triceps long head were fed through a HumBug noise eliminator. The remaining channels did not have additional filtering.

#### Experimental conditions

We tested 2 stimulation waveforms: i) the 2 ms conventional waveform, and ii) the 1 ms KHF waveform. Stimulation was applied using the same hardware as for lumbar tSCS. Paired pulses with an interstimulus interval of 33 ms were applied.

#### Cervical tSCS stimulation protocol

For each waveform, stimulation was first manually delivered by the experimenter to determine stimulation currents that evoke i) responses in the abductor pollicis brevis and the flexor carpi radialis at threshold, i.e., minimum stimulation currents to evoke responses with peak-to-peak amplitudes ≥ 50 µV in the two muscles, and ii) abductor pollicis brevis and flexor carpi radialis responses with maximum attainable peak-to-peak amplitudes. The stimulation currents then used in the cervical tSCS protocol were set to correspond to 50% and 100% of the current required to evoke responses at threshold as well as 5%, 25%, 50%, 75%, 100%, and 110% of the current required to evoke maximum responses in the two muscles.

Three paired pulses were applied at each stimulation amplitude, for a total of 48 repetitions per waveform. Pulses were delivered from lowest to highest stimulation current with 7±0.5 seconds (randomized) between repetitions. The testing order for the different waveforms was randomized (randarray, Microsoft Excel).

#### Data analysis and statistics

Data were analyzed and statistical comparisons were performed analogously to the lumbar tSCS protocol.

#### Peripheral conduction

Onset latencies of the responses in the abductor pollicis brevis were compared to the peripheral conduction time of this muscle. Peripheral conduction time was determined by applying 1 ms biphasic pulses to the median nerve at the wrist. A bipolar stimulating electrode (Digitimer E.SB020 Bipolar felt pad electrode) was placed between the flexor carpi radialis and the palmaris longus tendons and approximately 8 cm from the palmaris longus with the cathode distal to the anode^105^. Stimulation was incrementally increased from the M-wave threshold. At 125% of M_max_, 60 pulses were applied every 3 ± 0.5 seconds (randomized) to evoke F-waves^100–102^. Peripheral conduction time of the abductor pollicis brevis was then calculated using the same formula as for the medial gastrocnemius.

## Acknowledgements

R.K., L.L., R.H., and I.S received partial support from the National Institutes of Health NICHD Award Number K12HD073945, NINDS Award Number K01NS127936, and internal funding from the Department of Biomedical Engineering, the Department of Neurosurgery, and the McDonnell Center for Systems Neuroscience at Washington University in St. Louis. A.R. and Z.H. were funded by the German Federal Ministry of Education and Research (BMBF) project nr. 01ZZ2016. K.M. was supported by the Austrian Science Fund (FWF), project nr. P 34460-B.

## Author contributions

R.K., U.H., and Z.H., data analysis.

R.K., L.L., and N.B., software development.

R.K. and R.H., conducted experiments.

A.R., K.M, and I.S., conceptualization and supervision.

R.K., U.H., Z.H., R.H., A.R., K.M., and I.S., data interpretation.

R.K., U.H., A.R., K.M., and I.S., manuscript writing, review, and editing.

Z.H. and A.R., computational framework. Z.H., computational simulations.

## Data availability

Data from this study will be made available upon reasonable request to the corresponding author.

## Code availability

All software used to produce the figures in this manuscript will be available upon reasonable request to the corresponding author.

## Supplementary Data

**Supplementary Figure 1.**
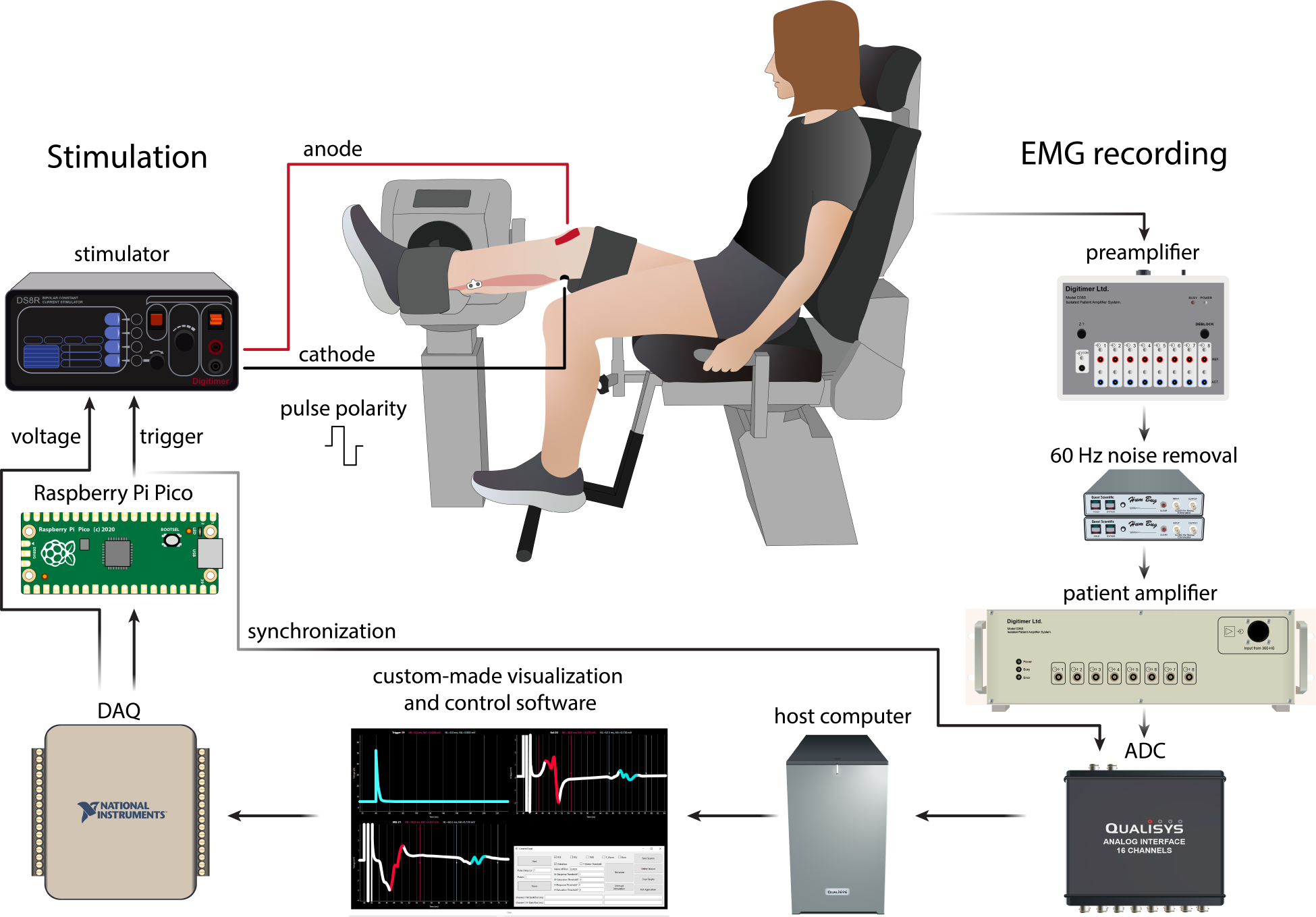
Technological platform for peripheral nerve stimulation experiments using a custom-built software. Wired bipolar EMG electrodes connected to a preamplifier (Digitimer D360, Digitimer Ltd, UK) were connected to a Hum Bug for removal of 60 Hz line noise before amplification (Hum Bug, Quest Scientific, India) and digitization (Qualysis Analog Interface 16 Channels, Qualisys, Sweden). Custom-built software (Python, v3.10.4) was used to record and display real-time EMG signals as well as segmented responses based on the TTL sync pulse from the constant current biphasic stimulator (Digitimer DS8R, Digitimer Ltd, UK). The software interface controlled a trigger pulse that was sent from the DAQ (USB-6001, National Instruments, USA) to the Raspberry Pi microcontroller which would in turn send a trigger to the stimulator (single trigger pulses in conventional waveforms, or multiple trigger pulses 100 µs apart in KHF waveforms). In addition, the DAQ sent a voltage signal to the stimulator to control the required stimulation current. The stimulator cathode (-) was connected to the popliteal fossa via a 3.2 cm diameter circular surface electrode and the 5 cm x 9 cm anode (+) was connected to the lower patella. The pulse polarity had a positive phase followed by a negative phase.

**Supplementary Figure 2.**
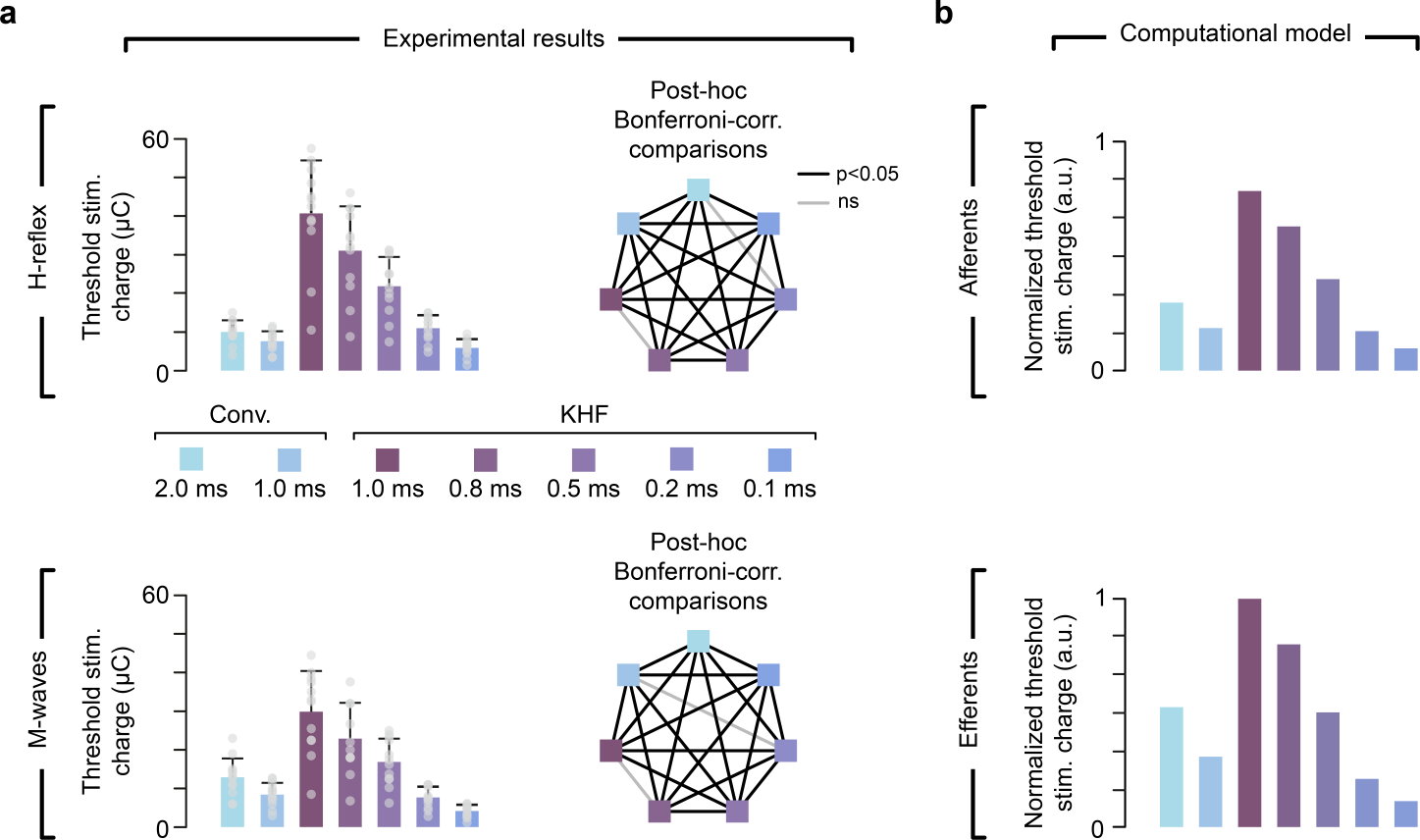
Effect of waveform and duration on H-reflex and M-wave charge at threshold. **a.** Stimulation charges required to evoke responses at threshold increased with increasing waveform duration for both the H-reflex and M-wave. Moreover, KHF waveforms with durations ≥ 0.5 ms required higher stimulation charge than conventional waveforms to elicit responses. **b.** Threshold stimulation charge required to depolarize afferent and efferent fibers in the computational model. Similarly to the experimental results, the threshold stimulation charge required to recruit a fiber increases with increasing burst duration. Note that the experimental setup measures EMG responses, while the computational model measures fiber recruitment. Bars represent estimated means while whiskers indicate the standard errors. The heptagon indicates the results for the post-hoc Bonferroni-corrected comparisons between waveforms; the solid black line indicates statistically significant differences between waveforms (p < 0.05). Abbreviations: stimulation (stim.), corrected (corr.), kilohertz frequency (KHF), conventional (Conv.), arbitrary units (a.u.), not significant (n.s.).

**Supplementary Figure 3.**
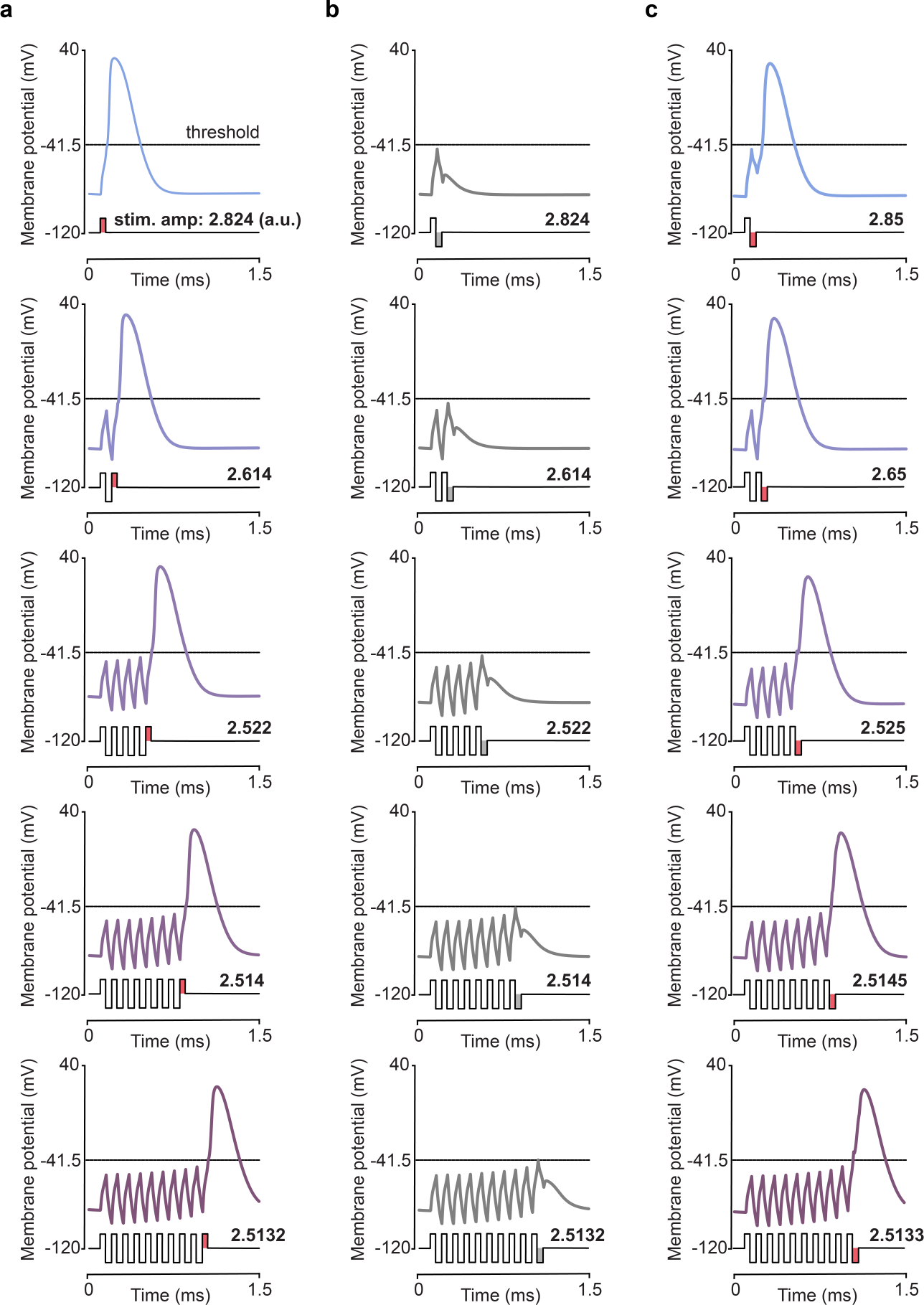
Negative phases in KHF waveforms counteract the generation of action potentials. **a.** Membrane potentials at threshold amplitudes in monophasic stimulation (top) and KHF waveforms of different durations ending in a positive phase. Stimulation amplitudes are shown above each waveform. At threshold amplitudes, action potentials are generated slightly after the last positive phase. **b**. Membrane potentials at the same stimulation amplitude as in **a** when an additional negative phase is added after the last positive phase. The negative phase prevents the action potential from occurring, so that threshold amplitudes that were eliciting an action potential in **a** can no longer do so. **c**. Membrane potentials at the stimulation amplitude necessary to elicit responses at threshold with the addition of the negative phase. Higher stimulation amplitudes are now required to elicit an action potential.

**Supplementary Figure 4.**
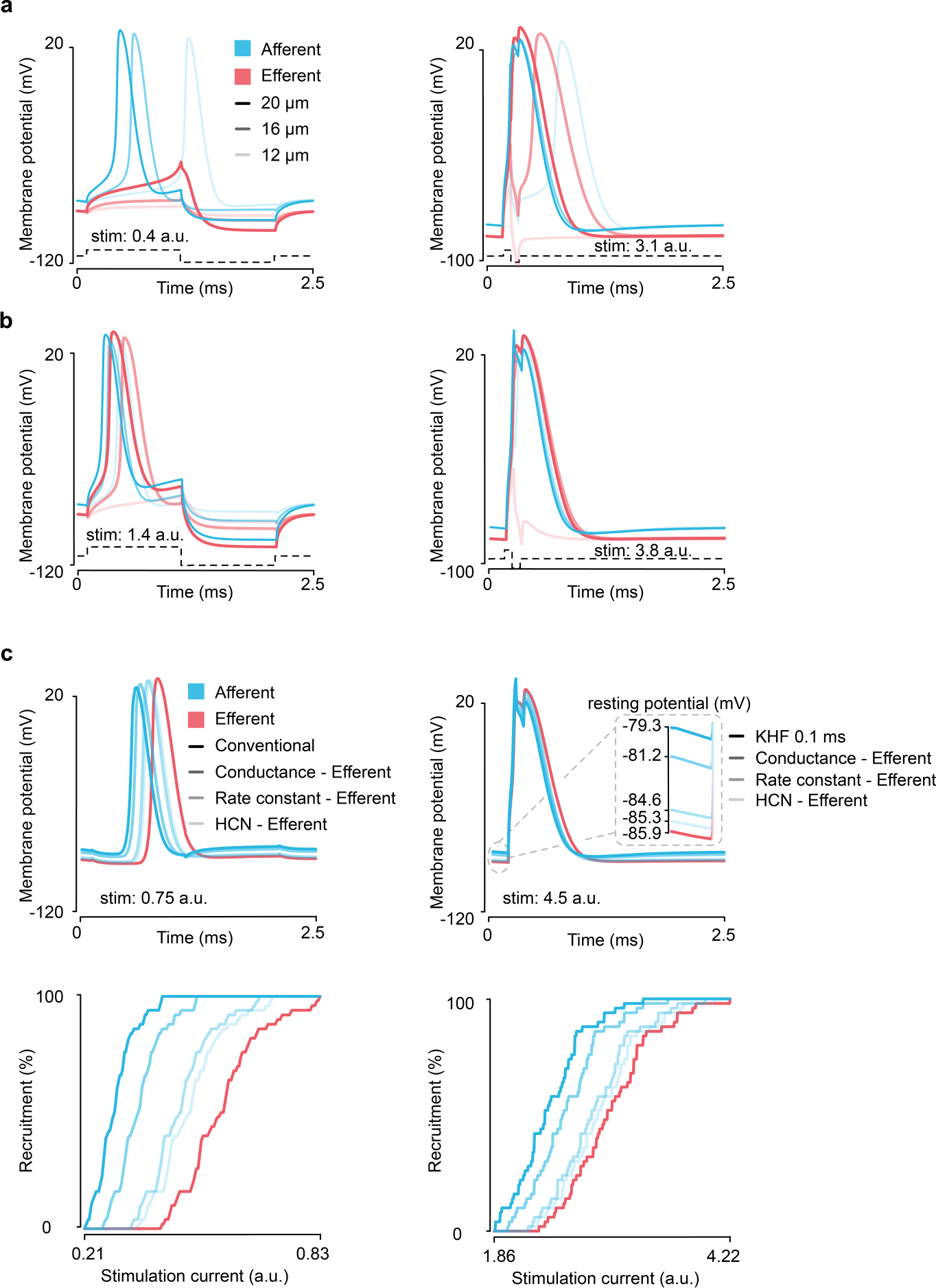
Effect of stimulation current and membrane parameters on recruitment of motor and proprioceptive fibers. **a.** Membrane potentials in fibers of different diameters during stimulation with 2 ms and 0.1 ms conventional pulses shown in Fig. 4 with a stimulation current of 0.4 a.u. **b.** Membrane potentials in fibers of different diameters with same stimulation waveforms as in **a** but with a higher stimulation current of 1.4 a.u. **c**. Membrane potentials and recruitment curves in efferent and afferent fibers when HCN-related membrane parameters in proprioceptive fibers were tuned to match those in efferent fibers at two stimulation currents, e.g., Conductance-Efferent: HCN parameters in afferent fibers related to conductance were tuned to match the HCN parameters in efferent fibers related to conductance. HCN-Efferent: all HCN-related parameters in afferent fibers were tuned to match those in efferent fibers. Note that changes in parameters related to the HCN channels, and hence their resting membrane potentials, were sufficient to explain differences in recruitment. Abbreviations: arbitrary units (a.u.), hyperpolarization-activated cyclic nucleotide-gated (HCN), kilohertz-frequency (KHF).

**Supplementary Figure 5.**
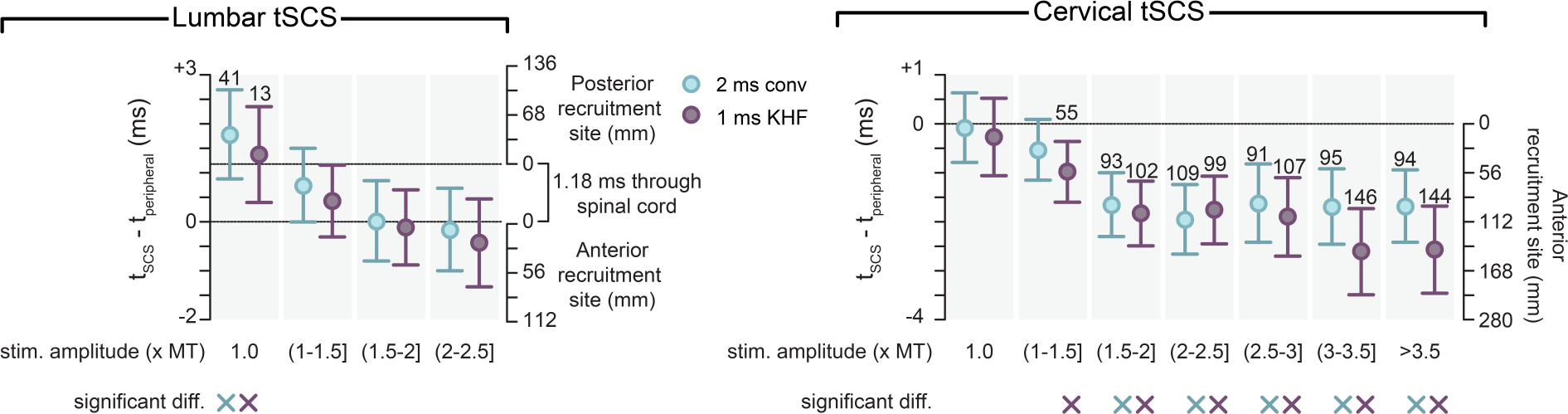
Action potential generation sites for lumbar and cervical tSCS. Action potential generation sites (recruitment sites) were estimated based on differences in latencies between tSCS and peripheral conduction times (as shown in Fig. 6). Conduction velocities were assumed to be 67.8 m/s for posterior roots and 55.8 m/s for anterior roots, while the transit time through the spinal cord (i.e., conduction time in afferent terminals plus synaptic delay) was assumed to be 1.18 ms^106,107^. Recruitment sites for responses that showed a significant difference between tSCS latencies and peripheral conduction times are shown above the whiskers. When the differences were not significant, recruitment sites were estimated to be within the respective roots at the closest end to the spinal cord. Central circles represent the estimated means while whiskers indicate the 95% confidence interval. The asterisks on the top of each muscle indicate the results for the post-hoc Bonferroni-corrected comparisons between waveforms; the × below the muscles indicate a statistically significant difference between t_tSCS_ and t_peripheral_.

**Supp. Table 1.**
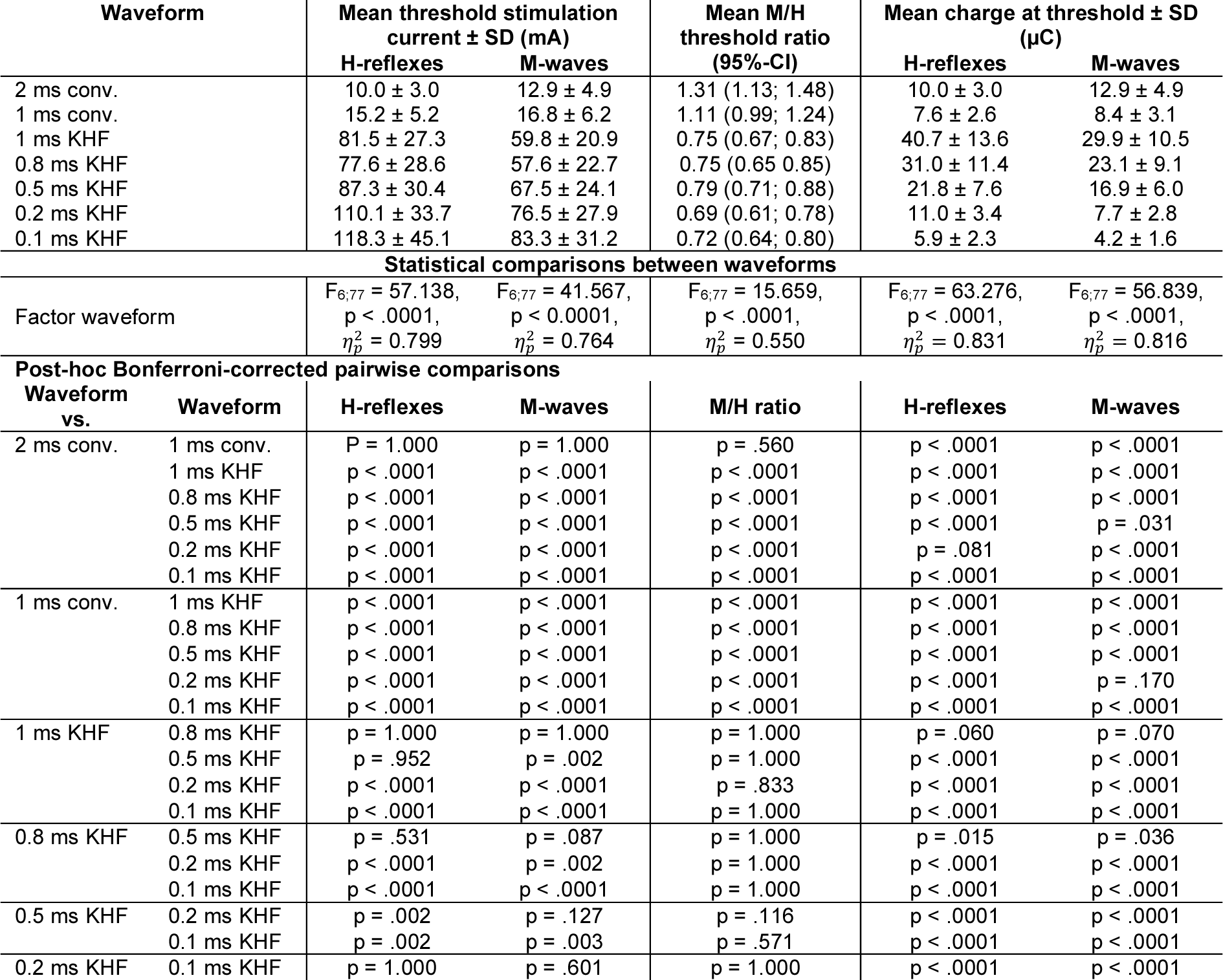
Threshold stimulation currents for eliciting H-reflexes and M-waves along with M/H threshold stimulation ratios and charges at threshold for the different waveforms tested. Threshold was defined as the minimum stimulation current required to elicit H-reflexes and M-waves, respectively, with peak-to-peak amplitudes ≥ 50 µV. Values are means ± standard deviation (SD) or 95%-confidence intervals (95%-CI) as indicated. Statistical comparisons were performed by fitting separate generalized linear mixed models with waveform as fixed factor and subject as a random factor. Abbreviations: conv., conventional waveform; KHF, kilohertz frequency waveform.

**Supp. Table 2.** Onset latencies of H-reflexes elicited with different waveforms as indicated at 1-1.5 times the respective threshold stimulation current. Values are means and 95%-confidence intervals (95%-CI). Statistical comparisons were performed by fitting a generalized linear mixed model with waveform as fixed factor and subject as a random factor. Abbreviations: conv., conventional waveform; KHF, kilohertz frequency waveform.

| Waveform |  | Mean onset latencies (ms) of H-reflexes at threshold (95%-CI) |
| --- | --- | --- |
| 2 ms conv. |  | 32.0 (30.3; 33.7) |
| 1 ms conv. |  | 31.5 (29.8; 33.1) |
| 1 ms KHF |  | 32.2 (30.7; 33.8) |
| 0.8 ms KHF |  | 31.9 (30.3;33.5) |
| 0.5 ms KHF |  | 31.7 (30.1; 33.3) |
| 0.2 ms KHF |  | 31.6 (30.0; 33.2) |
| 0.1 ms KHF |  | 31.2 (29.6; 32.9) |
| Statistical comparisons between waveforms |  |  |
| Factor waveform | | $F_{6;77} = 5.415$, $p = .0001$, $\eta_p^2 = 0.297$ |
| Post-hoc Bonferroni-corrected pairwise comparisons |  |  |
| Waveform vs. | Waveform |  |
| 2 ms conv. | 1 ms conv. | $P = 1.000$ |
| | 1 ms KHF | $p = 1.000$ |
| | 0.8 ms KHF | $p = 1.000$ |
| | 0.5 ms KHF | $p = 1.000$ |
| | 0.2 ms KHF | $p = 1.000$ |
| | 0.1 ms KHF | $p = .498$ |
| 1 ms conv. | 1 ms KHF | $p = .119$ |
| | 0.8 ms KHF | $p = 1.000$ |
| | 0.5 ms KHF | $p = 1.000$ |
| | 0.2 ms KHF | $p = 1.000$ |
| | 0.1 ms KHF | $p = 1.000$ |
| 1 ms KHF | 0.8 ms KHF | $p = .123$ |
| | 0.5 ms KHF | $p = .008$ |
| | 0.2 ms KHF | $p = .002$ |
| | 0.1 ms KHF | $p = .001$ |
| 0.8 ms KHF | 0.5 ms KHF | $p = 1.000$ |
| | 0.2 ms KHF | $p = .751$ |
| | 0.1 ms KHF | $p = .107$ |
| 0.5 ms KHF | 0.2 ms KHF | $p = 1.000$ |
| | 0.1 ms KHF | $p = .742$ |
| 0.2 ms KHF | 0.1 ms KHF | $p = 1.000$ |

**Supp. Table 3.** H_max_/M_max_ ratios along with peak-to-peak amplitudes of maximum H-reflexes (H_max_) and M-waves (M_max_) attained with the different waveforms tested. Values are means and 95%-confidence intervals (95%-CI). Statistical comparisons were performed by fitting separate generalized linear mixed models with waveform as fixed factor and subject as a random factor. Note that no post-hoc comparisons were performed for M_max_ since the statistical testing demonstrated no significant effect of the factor waveform on M_max_. Abbreviations: conv., conventional waveform; KHF, kilohertz frequency waveform; NA, not applicable.

| Waveform | Mean H<sub>max</sub>/M<sub>max</sub> ratio (95%-CI) | Mean H<sub>max</sub> (μV) (95%-CI) | Mean M<sub>max</sub> (μV) (95%-CI) |  |
| --- | --- | --- | --- | --- |
| 2 ms conv. | 0.54 (0.42; 0.66) | 3996.6 (2765.4; 5227.8) | 7624.2 (5396.7; 9851.8) |  |
| 1 ms conv. | 0.47 (0.35; 0.59) | 3456.1 (2459.2; 4453.0) | 7471.4 (5227.4; 9715.4) |  |
| 1 ms KHF | 0.38 (0.29; 0.47) | 2451.7 (1905.6; 2997.8) | 7287.6 (5048.4; 9526.7) |  |
| 0.8 ms KHF | 0.39 (0.29; 0.48) | 2478.9 (1937.8; 3019.9) | 7264.8 (5042.2; 9487.5) |  |
| 0.5 ms KHF | 0.39 (0.30; 0.48) | 2641.0 (2063.3; 3218.7) | 7412.6 (5189.7; 9635.4) |  |
| 0.2 ms KHF | 0.39 (0.30; 0.49) | 2451.5 (1849.7; 3053.3) | 7183.2 (4944.7; 9421.6) |  |
| 0.1 ms KHF | 0.34 (0.23; 0.45) | 2302.9 (1634.2; 2971.6) | 7304.2 (5065.3; 9543.2) |  |
| Statistical comparisons between waveforms |  |  |  |  |
| Factor waveform | F<sub>6;77</sub> = 3.365, p = .005, η<sub>p</sub><sup>2</sup> = 0.208 | F<sub>6;77</sub> = 2.595, p = .024, η<sub>p</sub><sup>2</sup> = 0.168 | F<sub>6;77</sub> = 1.530, p = .180, η<sub>p</sub><sup>2</sup> = 0.107 |  |
| Post-hoc Bonferroni-corrected pairwise comparisons |  |  |  |  |
| Waveform vs. | Waveform | H<sub>max</sub>/M<sub>max</sub> | H<sub>max</sub> | M<sub>max</sub> |
| 2 ms conv. | 1 ms conv. | p = 1.000 | p = 1.000 | NA |
|  | 1 ms KHF | p = .012 | p = .148 | NA |
|  | 0.8 ms KHF | p = .017 | p = .158 | NA |
|  | 0.5 ms KHF | p = .022 | p = .316 | NA |
|  | 0.2 ms KHF | p = .049 | p = .160 | NA |
|  | 0.1 ms KHF | p = .010 | p = .117 | NA |
| 1 ms conv. | 1 ms KHF | p = .285 | p = .325 | NA |
|  | 0.8 ms KHF | p = .390 | p = .347 | NA |
|  | 0.5 ms KHF | p = .497 | p = .804 | NA |
|  | 0.2 ms KHF | p = .766 | p = .356 | NA |
|  | 0.1 ms KHF | p = .163 | p = .280 | NA |
| 1 ms KHF | 0.8 ms KHF | p = 1.000 | p = 1.000 | NA |
|  | 0.5 ms KHF | p = 1.000 | p = 1.000 | NA |
|  | 0.2 ms KHF | p = 1.000 | p = 1.000 | NA |
|  | 0.1 ms KHF | p = 1.000 | p = 1.000 | NA |
| 0.8 ms KHF | 0.5 ms KHF | p = 1.000 | p = 1.000 | NA |
|  | 0.2 ms KHF | p = 1.000 | p = 1.000 | NA |
|  | 0.1 ms KHF | p = 1.000 | p = 1.000 | NA |
| 0.5 ms KHF | 0.2 ms KHF | p = 1.000 | p = 1.000 | NA |
|  | 0.1 ms KHF | p = 1.000 | p = 1.000 | NA |
| 0.2 ms KHF | 0.1 ms KHF | p = 1.000 | p = 1.000 | NA |

**Supp. Table 4.**
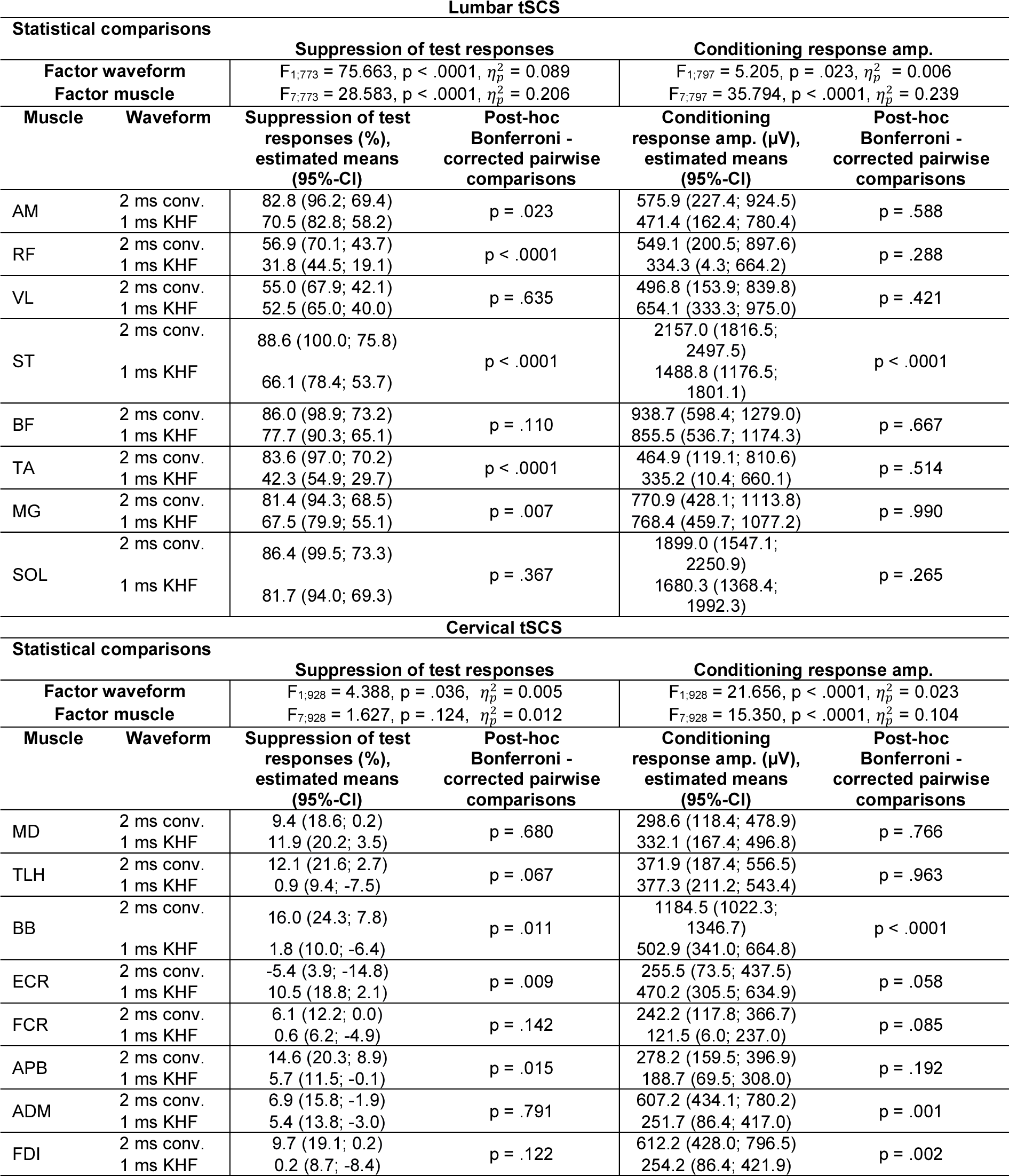
Post-activation depression of responses in lower and upper limb muscles evoked by 2 ms conventional (conv.) and 1 ms kilohertz frequency (KHF) waveforms applied through lumbar and cervical transcutaneous spinal cord stimulation (tSCS), respectively. Post-activation depression was tested by applying paired pulses with an interstimulus interval of 33 ms and calculating the ratio of the peak-to-peak amplitudes of the responses to the second (test response) and to the first stimuli (conditioning response)^17^. Suppression of test responses as well as peak-to-peak amplitudes of the conditioning responses were compared by fitting generalized linear mixed models with muscle and waveforms as fixed factors and subject as a random factor. Post-hoc Bonferroni-corrected pairwise comparisons were conducted between waveforms per muscle. Values are means and 95%-confidence intervals (95%-CI). Positive values indicate smaller test than conditioning responses, and positive values indicate larger test than conditioning responses. Abbreviations: ADM, abductor digiti minimi; AM, adductor magnus; APB, abductor pollicis brevis; BB, biceps brachii; BF, biceps femoris; ECR, extensor carpi radialis; FCR, flexor carpi radialis; FDI, first dorsal interosseus; MD, medial deltoid; MG, medial gastrocnemius; RF, rectus femoris; SOL, soleus; ST, semitendinosus; TA, tibialis anterior; TLH, triceps long head; VL, vastus lateralis.

**Supp. Table 5.** Comparison of post-activation depression of responses evoked by 2 ms conventional (conv.) and a ms kilohertz frequency (KHF) waveforms applied through lumbar vs. cervical transcutaneous spinal cord stimulation. Values are means and 95%-confidence intervals (95%-CI). Positive values indicate smaller test than conditioning responses. Statistical comparison was conducted by fitting a generalized linear mixed model with waveform and stimulation site as fixed factors and subject as a random factor.

| Statistical comparisons |  | Suppression of test responses (%), estimated means (95%-CI) |  |
| --- | --- | --- | --- |
| Factor waveform | | $F_{1;1729} = 56.059, p < .0001, \eta_p^2 = 0.031$ | |
| Factor stimulation site | | $F_{1; 1729} = 867.440, p < .0001, \eta_p^2 = 0.334$ | |
| Waveform | Stimulation site | Suppression of test responses (%),<br>estimated means (95%-CI) | Post-hoc Bonferroni -corrected<br>pairwise comparisons |
| 2 ms conv. | cervical | 10.8 (16.0; 5.7) | $p < .0001$ |
|  | lumbar | 76.0 (81.6; 70.4) |  |
| 1 ms KHF | cervical | 5.7 (10.7; 0.6) | $p < .0001$ |
|  | lumbar | 59.9 (65.3; 54.5) |  |

**Supp. Table 6.** Comparison of onset latencies of responses to 2 ms conventional (conv.) and 1 ms kilohertz frequency (KHF) waveforms applied through transcutaneous spinal cord stimulation (tSCS) and peripheral conduction times. For lumbar tSCS, responses in medial gastrocnemius, and for cervical tSCS, responses in abductor pollicis brevis were considered. Peripheral conduction times for these muscles were estimated based on their F-wave and M-wave latencies. Values are estimated mean differences between onset latencies and corresponding peripheral conduction times along with 95%-confidence intervals (95%-CI). Negative values indicate shorter onset latencies than peripheral conduction times, and positive values indicate longer onset latencies than peripheral conduction times. 95%-CIs with both the lower and upper limits assuming positive or negative values, respectively, indicate statistically significant differences (i.e., 0 not within the 95%-CI). Note that differences between onset latencies and peripheral conduction times were established for increasing stimulation currents. To this end, stimulation currents were binned as follows: bin 1, containing responses evoked at threshold; bin 2, (1-1.5] x threshold; bin 3, (1.5-2.0] x threshold; bin 4, (2.0-2.5] x threshold; bin 5, (2.5-3.0] x threshold; bin 6, (3.0-3.5] x threshold; and bin 7, >3.5 x threshold. For medial gastrocnemius, bins 1-4 were considered for further analysis, since no statistical difference in the number of available samples between conventional and KHF waveforms was found for these bins, χ2(3) = 3.162, p = .367, Cramer’s V = 0.133. For abductor pollicis brevis, all seven bins were considered, since no statistical difference in the number of available samples between conventional and KHF waveforms was found, χ2(6) = 4.218, p = .647, Cramer’s V = 0.104. Statistical comparisons were then performed by fitting generalized linear mixed models with waveform and stimulation current bin as fixed factors and subject as a random factor.

| Medial gastrocnemius |  |  |  |
| --- | --- | --- | --- |
| Statistical comparisons |  |  |  |
| Factor waveform | | $F_{1;147} = 2.616, p = .108, \eta_p^2 = 0.017$ | |
| Factor stimulation current bin | | $F_{3;147} = 18.832, p < .0001, \eta_p^2 = 0.278$ | |
| Bin | Waveform | $t_{\text{scs}} - t_{\text{peripheral}}$,<br>estimated means (95%-CI) | Post-hoc Bonferroni -corrected<br>pairwise<br>comparisons between waveforms |
| 1 | 2 ms conv.<br>1 ms KHF | 1.785 (0.875; 2.695)<br>1.372 (0.396; 2.348) | $p = .374$ |
| 2 | 2 ms conv.<br>1 ms KHF | 0.748 (-0.004; 1.500)<br>0.423 (-0.311; 1.158) | $p = .109$ |
| 3 | 2 ms conv.<br>1 ms KHF | -0.019 (-0.802; 0.763)<br>-0.115 (-0.883; 0.653) | $p = .713$ |
| 4 | 2 ms conv.<br>1 ms KHF | -0.158 (-1.002; 0.686)<br>-0.430 (-1.327; 0.466) | $p = .477$ |
| Abductor pollicis brevis |  |  |  |
| Statistical comparisons |  |  |  |
| Factor waveform | | $F_{1;360} = 10.216, p = .001, \eta_p^2 = 0.028$ | |
| Factor stimulation current bin | | $F_{6;360} = 23.016, p < .0001, \eta_p^2 = 0.277$ | |
| 1 | 2 ms conv.<br>1 ms KHF | -0.083 (-0.791; 0.625)<br>-0.267 (-1.058; 0.525) | $p = .583$ |
| 2 | 2 ms conv.<br>1 ms KHF | -0.534 (-1.151; 0.083)<br>-0.979 (-1.597; -0.361) | $p < .0001$ |
| 3 | 2 ms conv.<br>1 ms KHF | -1.652 (-2.298; -1.006)<br>-1.829 (-2.486; -1.173) | $p = .378$ |
| 4 | 2 ms conv.<br>1 ms KHF | -1.949 (-2.664; -1.233)<br>-1.759 (-2.446; -1.072) | $p = .476$ |
| 5 | 2 ms conv.<br>1 ms KHF | -1.621 (-2.421; -0.822)<br>-1.898 (-2.698; -1.097) | $p = .468$ |
| 6 | 2 ms conv.<br>1 ms KHF | -1.694 (-2.459; -0.928)<br>-2.607 (-3.491; -1.723) | $p = .027$ |
| 7 | 2 ms conv.<br>1 ms KHF | -1.682 (-2.416; -0.947)<br>-2.570 (-3.465; -1.675) | $p = .027$ |

**Supp. Table 7.** Post-activation depression of responses in medial gastrocnemius and abductor pollicis brevis evoked by 2 ms conventional and 1 ms kilohertz frequency (KHF) waveforms applied through lumbar and cervical transcutaneous spinal cord stimulation (tSCS), respectively. Post-activation depression was tested by applying paired pulses with an interstimulus interval of 33 ms and calculating the ratio of the peak-to-peak amplitudes of the responses to the second (test response) and to the first stimuli (conditioning response)^17^. Values are means and 95%-confidence intervals (95%-CI). Positive values indicate smaller test than conditioning responses, and negative values indicate larger test than conditioning responses. Note that levels of post-activation depression were established for incremental increases in stimulation currents. To this end, stimulation currents were binned as follows: bin 1, containing responses evoked at threshold; bin 2, (1-1.5] x threshold; bin 3, (1.5-2.0] x threshold; bin 4, (2.0-2.5] x threshold; bin 5, (2.5-3.0] x threshold; bin 6, (3.0-3.5] x threshold; and bin 7, >3.5 x threshold. For medial gastrocnemius, bins 1-4 were considered for further analysis, since no statistical difference in the number of available samples between conventional and KHF waveforms was found for these bins, χ2(3) = 3.162, p = .367, Cramer’s V = 0.133. For abductor pollicis brevis, all seven bins were considered, since no statistical difference in the number of available samples between conventional and KHF waveforms was found, χ2(6) = 4.218, p = .647, Cramer’s V = 0.104. Statistical comparisons were then performed by fitting generalized linear mixed models with waveform and stimulation current bin as fixed factors and subject as a random factor.

| Medial gastrocnemius |  |  |  |  |
| --- | --- | --- | --- | --- |
| Statistical comparisons |  |  |  |  |
| Factor waveform | | $F_{1;166} = 19.163, p < .0001, \eta_p^2 = 0.103$ | | |
| Factor stimulation current bin | | $F_{3;166} = 9.090, p < .0001, \eta_p^2 = 0.141$ | | |
| Suppression of test responses (%) ± SE | | | | |
| Bin |  | 2 ms conventional waveform | 1 ms KHF waveform |  |
| 1 | | 72.7 ± 7.3 | 46.6 ± 7.8 | |
| 2 | | 83.8 ± 5.8 | 71.6 ± 5.5 | |
| 3 | | 91.1 ± 6.1 | 75.9 ± 6.1 | |
| 4 | | 87.3 ± 7.2 | 81.7 ± 7.9 | |
| Bin vs. Bin |  | Post-hoc Bonferroni -corrected pairwise comparisons |  |  |
| 1 | 2 | $p = .326$ | $p = .001$ | |
| | 3 | $p = .036$ | $p < .0001$ | |
| | 4 | $p = .289$ | $p < .0001$ | |
| 2 | 3 | $p = .397$ | $p = .693$ | |
| | 4 | $p = 1.000$ | $p = .404$ | |
| 3 | 4 | $p = 1.000$ | $p = .693$ | |
| Abductor pollicis brevis |  |  |  |  |
| Statistical comparisons |  |  |  |  |
| Factor waveform | | $F_{1;375} = 0.389, p = .533, \eta_p^2 = 0.001$ | | |
| Factor stimulation current bin | | $F_{6;375} = 0.578, p = .748, \eta_p^2 = 0.009$ | | |
| Bin |  | 2 ms conventional waveform | 1 ms KHF waveform |  |
| 1 | | 9.3 ± 7.6 | 10.4 ± 7.8 | |
| 2 | | 14.9 ± 3.7 | 4.2 ± 3.7 | |
| 3 | | 9.1 ± 5.3 | 3.9 ± 5.6 | |
| 4 | | 2.6 ± 7.9 | 6.9 ± 6.9 | |
| 5 | | 0.4 ± 10.4 | -1.0 ± 10.5 | |
| 6 | | 9.4 ± 9.5 | 2.7 ± 12.7 | |
| 7 | | -1.1 ± 8.5 | -1.9 ± 12.3 | |

**Supp. Table 8.**
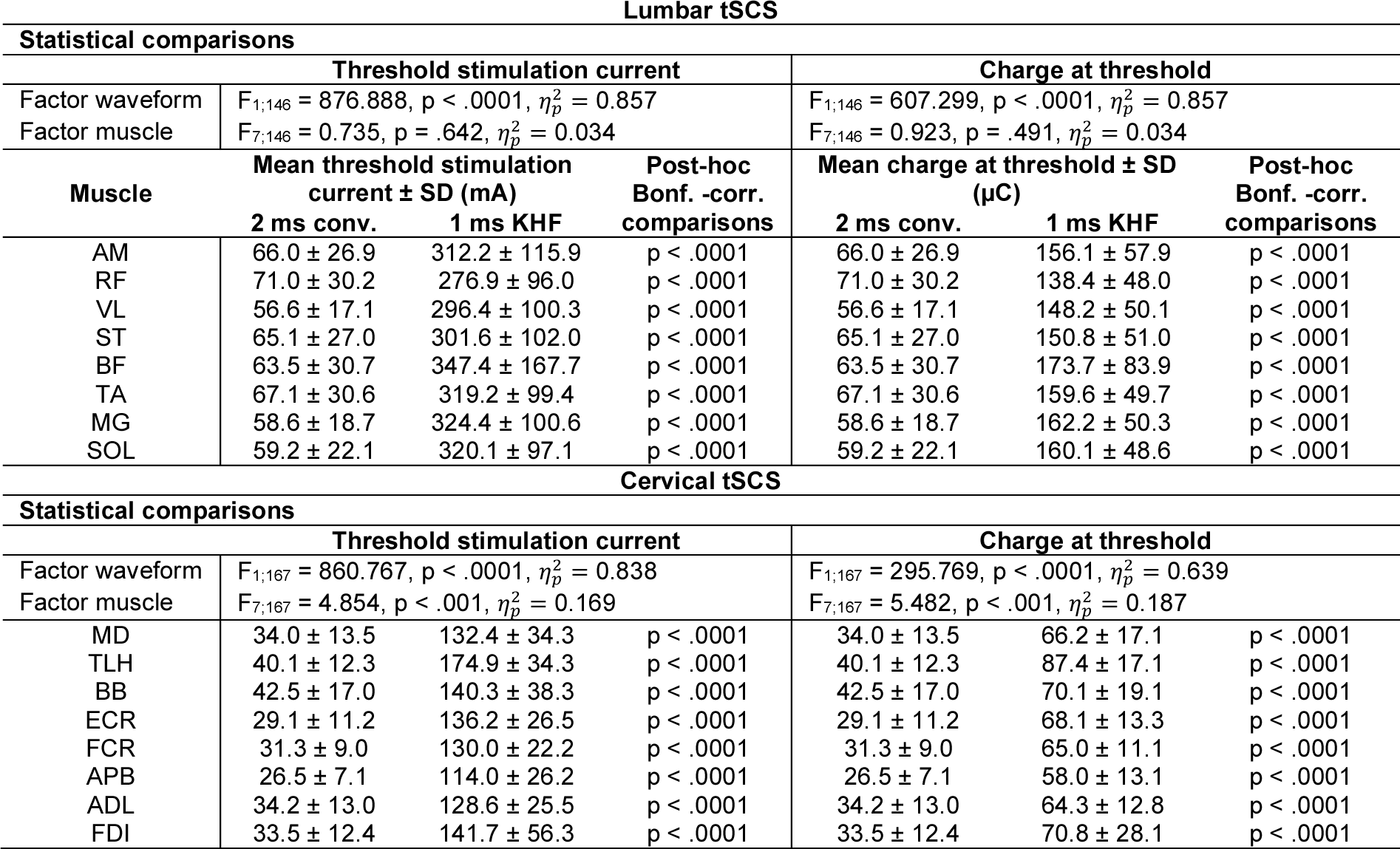
Threshold stimulation currents and charges at threshold for eliciting responses in lower limb and upper limb muscles evoked by 2 ms conventional and 1 ms kilohertz frequency (KHF) waveforms applied through lumbar and cervical transcutaneous spinal cord stimulation (tSCS), respectively. Threshold was defined as the minimum stimulation current required to elicit responses with peak-to-peak amplitudes ≥ 50 µV. Values are means ± the standard deviation (SD). Statistical comparisons were performed by fitting generalized linear mixed models with waveform and muscle as fixed factors and subject as random factor. Abbreviations: ADM, abductor digiti minimi; AM, adductor magnus; APB, abductor pollicis brevis; BB, biceps brachii; BF, biceps femoris; ECR, extensor carpi radialis; FCR, flexor carpi radialis; FDI, first dorsal interosseus; MD, medial deltoid; MG, medial gastrocnemius; RF, rectus femoris; SOL, soleus; ST, semitendinosus; TA, tibialis anterior; TLH, triceps long head; VL, vastus lateralis.

Lumbar tSCS
| Statistical comparisons |  |  |  |  |  |  |
| --- | --- | --- | --- | --- | --- | --- |
| Threshold stimulation current |  |  |  | Charge at threshold |  |  |
| Factor waveform | $F_{1;146} = 876.888, p < .0001, \eta_p^2 = 0.857$ | | | $F_{1;146} = 607.299, p < .0001, \eta_p^2 = 0.857$ | | |
| Factor muscle | $F_{7;146} = 0.735, p = .642, \eta_p^2 = 0.034$ | | | $F_{7;146} = 0.923, p = .491, \eta_p^2 = 0.034$ | | |
| Muscle | Mean threshold stimulation current ± SD (mA) | | Post-hoc Bonf. -corr. comparisons | Mean charge at threshold ± SD (μC) | | Post-hoc Bonf. -corr. comparisons |
|  | 2 ms conv. | 1 ms KHF |  | 2 ms conv. | 1 ms KHF |  |
| AM | 66.0 ± 26.9 | 312.2 ± 115.9 | $p < .0001$ | 66.0 ± 26.9 | 156.1 ± 57.9 | $p < .0001$ |
| RF | 71.0 ± 30.2 | 276.9 ± 96.0 | $p < .0001$ | 71.0 ± 30.2 | 138.4 ± 48.0 | $p < .0001$ |
| VL | 56.6 ± 17.1 | 296.4 ± 100.3 | $p < .0001$ | 56.6 ± 17.1 | 148.2 ± 50.1 | $p < .0001$ |
| ST | 65.1 ± 27.0 | 301.6 ± 102.0 | $p < .0001$ | 65.1 ± 27.0 | 150.8 ± 51.0 | $p < .0001$ |
| BF | 63.5 ± 30.7 | 347.4 ± 167.7 | $p < .0001$ | 63.5 ± 30.7 | 173.7 ± 83.9 | $p < .0001$ |
| TA | 67.1 ± 30.6 | 319.2 ± 99.4 | $p < .0001$ | 67.1 ± 30.6 | 159.6 ± 49.7 | $p < .0001$ |
| MG | 58.6 ± 18.7 | 324.4 ± 100.6 | $p < .0001$ | 58.6 ± 18.7 | 162.2 ± 50.3 | $p < .0001$ |
| SOL | 59.2 ± 22.1 | 320.1 ± 97.1 | $p < .0001$ | 59.2 ± 22.1 | 160.1 ± 48.6 | $p < .0001$ |

**Supp. Table 8.** Threshold stimulation currents and charges at threshold for eliciting responses in lower limb and upper limb muscles evoked by 2 ms conventional and 1 ms kilohertz frequency (KHF) waveforms applied through lumbar and cervical transcutaneous spinal cord stimulation (tSCS), respectively.
| Statistical comparisons |  |  |  |  |  |  |
| --- | --- | --- | --- | --- | --- | --- |
| Threshold stimulation current |  |  |  | Charge at threshold |  |  |
| Factor waveform | $F_{1;167} = 860.767, p < .0001, \eta_p^2 = 0.838$ | | | $F_{1;167} = 295.769, p < .0001, \eta_p^2 = 0.639$ | | |
| Factor muscle | $F_{7;167} = 4.854, p < .001, \eta_p^2 = 0.169$ | | | $F_{7;167} = 5.482, p < .001, \eta_p^2 = 0.187$ | | |
| | Mean threshold stimulation current ± SD (mA) | | Post-hoc Bonf. -corr. comparisons | Mean charge at threshold ± SD (μC) | | Post-hoc Bonf. -corr. comparisons |
|  | 2 ms conv. | 1 ms KHF |  | 2 ms conv. | 1 ms KHF |  |
| MD | 34.0 ± 13.5 | 132.4 ± 34.3 | $p < .0001$ | 34.0 ± 13.5 | 66.2 ± 17.1 | $p < .0001$ |
| TLH | 40.1 ± 12.3 | 174.9 ± 34.3 | $p < .0001$ | 40.1 ± 12.3 | 87.4 ± 17.1 | $p < .0001$ |
| BB | 42.5 ± 17.0 | 140.3 ± 38.3 | $p < .0001$ | 42.5 ± 17.0 | 70.1 ± 19.1 | $p < .0001$ |
| ECR | 29.1 ± 11.2 | 136.2 ± 26.5 | $p < .0001$ | 29.1 ± 11.2 | 68.1 ± 13.3 | $p < .0001$ |
| FCR | 31.3 ± 9.0 | 130.0 ± 22.2 | $p < .0001$ | 31.3 ± 9.0 | 65.0 ± 11.1 | $p < .0001$ |
| APB | 26.5 ± 7.1 | 114.0 ± 26.2 | $p < .0001$ | 26.5 ± 7.1 | 58.0 ± 13.1 | $p < .0001$ |
| ADL | 34.2 ± 13.0 | 128.6 ± 25.5 | $p < .0001$ | 34.2 ± 13.0 | 64.3 ± 12.8 | $p < .0001$ |
| FDI | 33.5 ± 12.4 | 141.7 ± 56.3 | $p < .0001$ | 33.5 ± 12.4 | 70.8 ± 28.1 | $p < .0001$ |

**Supp Table 9.** Participant demographics information for the neurophysiology experiments.

| ID | Age (y) | Sex | Height (m) | Weight (kg) | Race | Hispanic/Latino | Experiment |  |  |
| --- | --- | --- | --- | --- | --- | --- | --- | --- | --- |
|  |  |  |  |  |  |  | Peripheral | Cervical (Protocol 2) | Lumbar |
| RS001 | 35 | Male | 1.67 | 60 | White | Yes | ✓ |  | ✓ |
| RS002 | 27 | Female | 1.7 | 61 | White | No | ✓ |  | ✓ |
| RS004 | 29 | Male | 1.85 | 70 | White | No | Monophasic stimulation |  |  |
| RS005 | 23 | Male | 1.85 | 84 | White | No | Monophasic stimulation | ✓ |  |
| RS006 | 23 | Female | 1.63 | 50 | White | No | Monophasic stimulation |  |  |
| RS007 | 22 | Female | 1.57 | 45 | Asian | No | Incomplete dataset |  |  |
| RS008 | 22 | Female | 1.78 | 59 | White | No | ✓ |  |  |
| RS009 | 23 | Female | 1.83 | 79 | White | No | ✓ |  | ✓ |
| RS010 | 27 | Male | 1.83 | 88 | White | No | ✓ |  |  |
| RS011 | 22 | Female | 1.65 | 54 | White | No | ✓ |  | ✓ |
| RS012 | 25 | Male | 1.63 | 59 | Asian | No | ✓ | ✓ | ✓ |
| RS013 | 22 | Female | 1.65 | 59 | White | Yes | ✓ | ✓ |  |
| RS014 | 25 | Preferred not to identify | 1.85 | 75 | Asian | No | ✓ | ✓ |  |
| RS015 | 28 | Male | 1.85 | 84 | Asian, White | No | Monophasic stimulation | ✓ | ✓ |
| RS016 | 28 | Female | 1.75 | 64 | White | No | ✓ |  | ✓ |
| RS017 | 25 | Female | 1.69 | 64 | White | No |  | ✓ |  |
| RS018 | 23 | Female | 1.68 | 66 | Asian, White | No |  | ✓ |  |
| RS019 | 24 | Male | 1.78 | 64 | White | No |  | ✓ |  |
| RS020 | 23 | Female | 1.63 | 73 | White | No |  | ✓ |  |
| RS021 | 24 | Female | 1.64 | 52 | Middle Eastern | No | ✓ |  |  |
| RS022 | 27 | Female | 1.72 | 65 | Asian | No | ✓ |  |  |
| RS023 | 25 | Female | 1.65 | 60 | White | No |  | ✓ |  |
| RS024 | 28 | Male | 1.7 | 82 | White | No |  |  | ✓ |
| RS025 | 28 | Female | 1.75 | 68 | White, American Indian or Alaska Native | Yes |  |  | ✓ |
| RS026 | 23 | Male | 1.73 | 61 | White | No |  |  | ✓ |

